# Genome wide identification of NAC transcription factors and their role in abiotic stress tolerance in *Chenopodium quinoa*

**DOI:** 10.1101/693093

**Authors:** Nouf Owdah Alshareef, Elodie Rey, Holly Khoury, Mark Tester, Sandra M. Schmöckel

## Abstract

*Chenopodium quinoa* Willd. (quinoa) is a pseudocereal with high nutritional value and relatively high tolerance to several abiotic stresses, including water deficiency and salt stress, making it a suitable plant for the study of mechanisms of abiotic stress tolerance. NAC (NAM, ATAF and CUC) transcription factors are involved in a range of plant developmental processes and in the response of plants to biotic and abiotic stresses. In the present study, we perform a genome-wide comprehensive analysis of the *NAC* transcription factor gene family in quinoa. In total, we identified 107 quinoa *NAC* transcription factor genes, distributed equally between sub-genomes A and B. They are phylogenetically clustered into two major groups and 18 subgroups. Almost 75% of the identified *CqNAC* genes were duplicated two to seven times and the remaining 25% of the *CqNAC* genes were found as a single copy. We analysed the transcriptional responses of the identified quinoa *NAC* TF genes in response to various abiotic stresses. The transcriptomic data revealed 28 stress responsive *CqNAC* genes, where their expression significantly changed in response to one or more abiotic stresses, including salt, water deficiency, heat and phosphate starvation. Among these stress responsive NACs, some were previously known to be stress responsive in other species, indicating their potentially conserved function in response to abiotic stress across plant species. Six genes were differentially expressed specifically in response to phosphate starvation but not to other stresses, and these genes may play a role in controlling plant responses to phosphate deficiency. These results provide insights into quinoa *NACs* that could be used in the future for genetic engineering or molecular breeding.

## 1. Introduction

The NAC transcription factor (TF) family is plant specific and is one of the largest families of TFs in plants. The acronym for NAC is derived from three TFs: NAM, ATAF and CUC, where NAM is an acronym for No Apical Meristem (1), ATAF stands for Arabidopsis Transcription Activator Factor (2), and CUC is a Cup Shaped Cotyledon (1). These three genes share a conserved N-terminal DNA-binding NAC domain. *Arabidopsis thaliana* (Arabidopsis) has 117 NACs, also called ANACs (3), rice has 151 NACs called ONACs (3), and soybean has 152 NACs (4). NAC proteins regulate a wide range of physiological and developmental processes – for example, petunia NAM and Arabidopsis CUC1-2 proteins are involved in shoot meristem development (1, 2). Other NAC members are involved in floral morphology (1, 5), plant senescence (6, 7), cell division (8), cell wall synthesis (9) and lateral root development (10).

Typical NAC TF proteins are characterized by a highly conserved DNA binding NAC domain at the N-terminus region. The NAC domain spans approximately 150 amino acids and consists of five conserved subdomains (A-E) that make up motifs for DNA binding, protein-protein interaction or transcription factor dimerization (11). Subdomains C and D are conserved and bind to DNA, subdomain A is involved in NAC dimerization, and subdomains B and E are highly divergent and may contribute to the functional diversity of NAC TFs (11, 12). The C-terminal region of NAC TFs is diversified and contains the transcription regulatory domain, which contributes to either transcription activation or repression (11, 13, 14).

Some NAC TF proteins possess transmembrane motifs in the C-terminal region that anchor NAC proteins to the intracellular membranes and make it inactive (15). When the NAC protein is activated, it undergoes proteolytic cleavage to release the NAC protein from the membrane to enable its TF function in the nucleus. These membrane-associated NAC TFs are designated as NAC with transmembrane motif1-like (NTL); most of them are associated with plasma membranes and a few are anchored to the endoplasmic reticulum membrane (15). The size of these NTL TFs is larger (from 335-652 amino acids) than the non-membrane associated NAC TFs, which are usually around 320 residues (15). More than 13 NAC TFs in Arabidopsis and six in rice have been described as NTL (15).

Some variation in the structure of NAC proteins has been reported. These NACs are called atypical NAC proteins or NAC-like proteins. These variant proteins include some proteins that have only a NAC domain without a C-terminal region (16, 17) and other proteins that have a tandem repeat of the NAC domains (12). Another two variants of NAC proteins are: suppressor of gamma response 1 (SOG1) proteins that have an extra sequence preceding the conserved NAC domain (17, 18), and vascular plant one-zinc-finger (VOZ) proteins that have a DNA-binding zinc finger, transcriptional regulation domain (TRD) at the N-terminal and a NAC domain at the C-terminus (12–14, 19).

Quinoa (*Chenopodium quinoa Willd.*, 2n=4x=36) is a dicotyledonous allotetraploid pseudocereal plant with 18 chromosomes (20). It belongs to the Amaranthaceae family, which also includes other economically important crops such as *Beta vulgaris* (beet), *Spinacia oleracea* (spinach) and *Amaranthus hypochondriacus* (amaranth). Quinoa has recently gained much attention due to its high nutritional value and high tolerance to environmental stresses. The grain of quinoa has a balanced ratio of carbohydrates, lipids and protein, a higher content of essential amino acids and is rich in iron and vitamins such as vitamin B1, B6 and E (21). In addition to quinoa’s nutritional value, it has a high tolerance to different abiotic stresses, including low temperature, drought and salinity (22, 23). Quinoa maintains the highest biomass when grown at 100 mM NaCl and the biomass is reduced by up to 50% when it grows under 500 mM NaCl (24). These traits make quinoa a good model for understanding the mechanisms of stress tolerance.

The recent completion of the genome of quinoa has allowed to study NAC genes at the whole genome level. To date, only the HSP17 and WRKY gene families have been systematically analysed in quinoa (25, 26). Here, we identify 107 NAC TF genes in the quinoa genome and investigate their transcriptional responses to different stresses including salt, drought and heat. We perform a systematic analysis of NAC TFs in quinoa using the available high-quality reference genome sequence of quinoa (27). This study provides a basis for future functional characterization of NAC TFs in quinoa that could be used in quinoa stress-tolerance research.

## 2. Results and discussion

### 2.1 Identification of the CqNAC transcription factors family in quinoa

In order to identify NAC TFs in the genome of quinoa, we used a combination of several search methods that have been used previously to identify NACs in different plant species. We employed especially those methods that identified NACs in species with duplicated genomes, such as *Panicum virgatum* (Switchgrass), *Populus trichocarpa* (black cottonwood) and *Gossypium raimondii* (cotton) (28–31).

We used the reference genome of quinoa accession QQ74 (27) to identify NAC TFs in quinoa. First, we used the Hidden Markov Model (HMM) profile of the protein family (Pfam) “NAC domain” (PF02365) as a query to search for proteins containing NAC domains in the genome of quinoa using the phytozome database version12 (phytozome, http://www.phytozome.net, version 12). This search identified 104 putative NACs proteins. Then, we employed a basic local alignment tool for proteins (BLASTp) against QQ74 peptides using the NAC domain sequences of the 110 Arabidopsis NACs. This search identified 109 putative NACs (six more than the Pfam search using the phytozome database). To find further NACs in the genome of quinoa, we used the TF prediction tool from the Plant Transcription Factor Database (PTFDB, http://planttfdb.cbi.pku.edu.cn/, version 4.0) (32) to predict all TFs in the peptide sequences of quinoa as an input (44,776 peptide sequences). We were able to identify 2093 TFs belonging to different families (S1 Table); among those, 107 CqTFs belong to the NAC family. As a quality control and to verify the reliability of our search, we ran the same search method for Arabidopsis and verified that we received the expected number of Arabidopsis NACs.

In order to identify the total number of NAC TFs in quinoa, we generated a combined list of the three search approaches (Pfam, BLASTp and PTFDB) (Fig 1, S2 Table). This list consisted of 112 putative CqNACs. All of the three search methods resulted in a similar number of CqNACs (Fig 1). There were 103 CqNACs common to all of the search methods, with two CqNACs identified only in the PTFDB search and five CqNACs identified in the BLASTp search only. To further confirm these putative CqNAC genes, we scanned the protein sequence of all 112 putative CqNAC proteins for the presence of the NAC domain using the InterProScan program (http://www.ebi.ac.uk/Tools/InterProScan/), the hmmscan function of the HMMER web server (https://www.ebi.ac.uk/Tools/hmmer/search/hmmscan, HmmerWeb version 2.28.0) (33) and conserved domain search (CDS) of the NCBI database. The presence of the NAC domain was confirmed in 107 CqNACs sequences, whereas the NAC domain was not present in the remaining five putative CqNAC proteins (all had been identified from the BLASTp search) (Fig 1, S2 Table). All of these search methods suggest that only 107 CqNACs genes exist in the reference genome of quinoa. The confirmed 107 putative quinoa NACs are listed in Table 1 and are used for further analyses.

**Fig 1.**
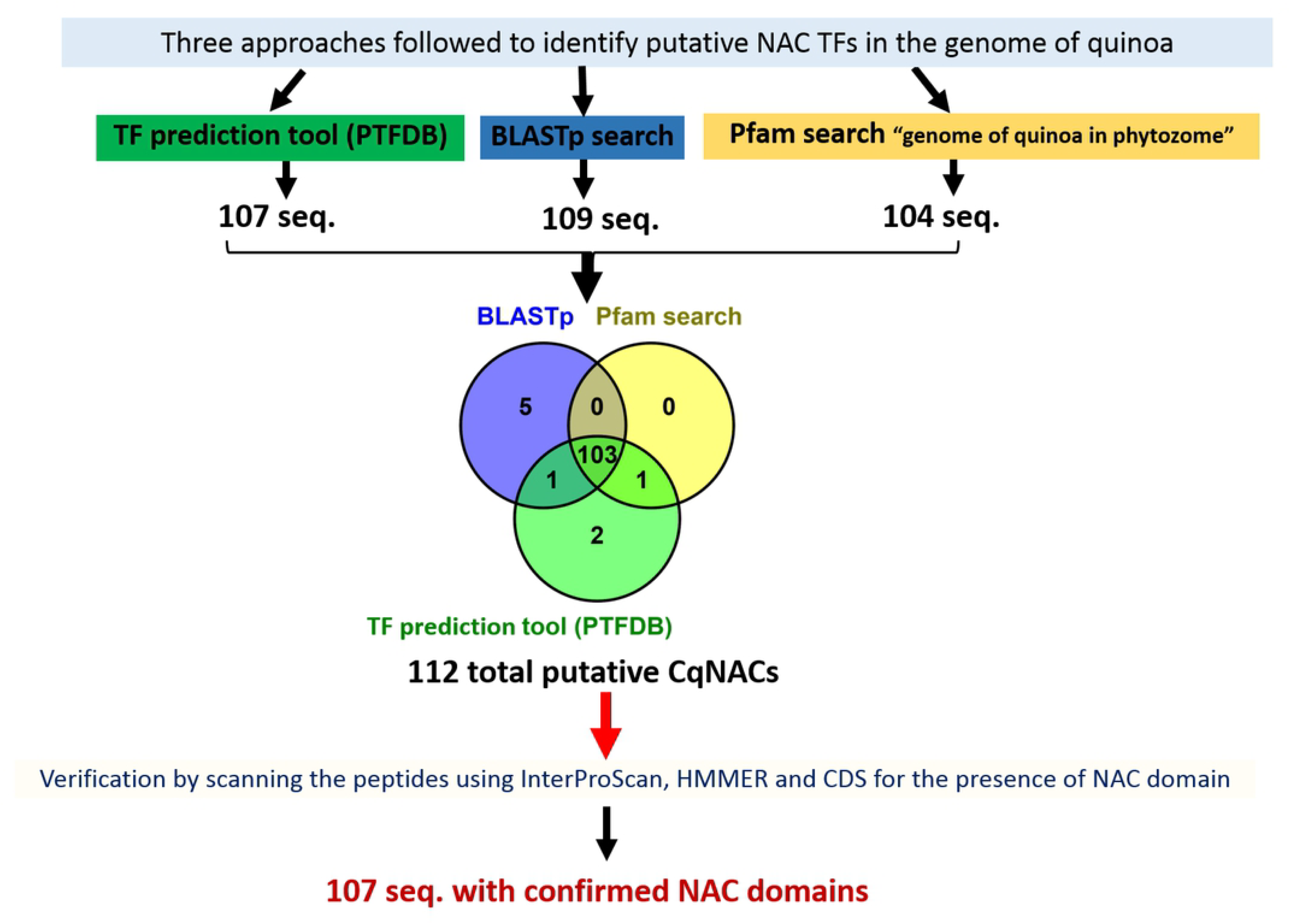
Scheme representing the bioinformatic approaches used to identify NAC genes in the genome of quinoa. A combined list of three search approaches was generated (using Pfam, BLASTp and the TF prediction tool of the Plant Transcription Factor Database (PTFDB)). Scanned of all 112 putative CqNAC proteins for the presence of the NAC domain using, the hmmscan function of the HMMER web server and conserved domain search (CDS) of the NCBI database. A total of 107 CqNAC proteins were identified.

**Table 1.** NAC transcription factor family in quinoa. Illustrating the sub-genome location, start and end of each gene, gene length, number of exons and protein length. CqNACs in bold with (*) sign are membrane associated CqNAC predicted by TMHMM server v.2.0. (http://www.cbs.dtu.dk/services/TMHMM/).

| Nb. | Gene ID | Gene Name | Brief Description | chromosomal location_subgenome | START | END | gene length (bp) | Nb. exons | protein size (aa) |
| --- | --- | --- | --- | --- | --- | --- | --- | --- | --- |
| 1 | AUR62017896 | CqNAC02 | NAC002:NACdomain-containingprotein2 (ATAF-1) | cq15_A | 20376810 | 20378712 | 1903 | 3 | 224 |
| 2 | AUR62003583 | CqNAC02 | NAC002:NACdomain-containingprotein2 (ATAF-1) | cq9_B | 3900006 | 3901944 | 1939 | 3 | 224 |
| 3 | AUR62000536 | CqNAC02 | NAC002:NACdomain-containingprotein2 (ATAF-1) | cq12_A | 6131839 | 6134224 | 2386 | 4 | 248 |
| 4 | AUR62015520 | CqNAC07 | NAC007: NAC domain-containing protein 7 | cq11_B | 21032546 | 21034478 | 1933 | 4 | 248 |
| 5 | AUR62002191 | CqNAC07 | NAC007: NAC domain-containing protein 7 | cq7_A | 65990389 | 65992323 | 1935 | 4 | 249 |
| 6 | AUR62007469 | CqNAC07 | NAC007: NAC domain-containing protein 7 | cq13_A | 728734 | 730378 | 1645 | 3 | 370 |
| 7 | AUR62039927 | CqNAC07 | NAC007: NAC domain-containing protein 7 | cq16_B | 78627158 | 78632662 | 5505 | 4 | 370 |
| 8 | AUR62004572 | CqNAC08 | NAC008: NAC domain-containing protein 8 | cq1_B | 120553744 | 120556805 | 3062 | 6 | 388 |
| 9 | AUR62022685 | CqNAC08 | NAC008: NAC domain-containing protein 8 | cq10_A | 5677119 | 5680213 | 3095 | 6 | 374 |
| 10 | AUR62002922 | CqNAC08 | NAC008: NAC domain-containing protein 8 | cq6_B | 15831647 | 15834922 | 3276 | 7 | 339 |
| 11 | AUR62035283 | CqNAC08 | NAC008: NAC domain-containing protein 8 | cq14_A | 9028304 | 9032559 | 4256 | 7 | 365 |
| 12 | AUR62029330 | CqNAC10 | NAC010: NAC domain-containing protein 10 | cq12_A | 51513524 | 51515901 | 2378 | 5 | 328 |
| 13 | AUR62031305 | CqONAC010 | ONAC010: NAC transcription factor ONAC010 | cq1_B | 25489514 | 25491226 | 1713 | 2 | 158 |
| 14 | AUR62027390 | CqONAC010 | ONAC010: NAC transcription factor ONAC010 | cq2_A | 40025813 | 40027566 | 1754 | 2 | 168 |
| 15 | AUR62016213 | CqONAC010 | ONAC010: NAC transcription factor ONAC010 | cq7_A | 75752855 | 75754429 | 1575 | 3 | 389 |
| 16 | AUR62011638 | CqONAC010 | ONAC010: NAC transcription factor ONAC010 | cq15_A | 13499168 | 13500708 | 1541 | 3 | 414 |
| 17 | **AUR62033952** | **CqNAC17** | **NAC017: NAC domain-containing protein 17 \*** | **cq0_** | **184747668** | **184752783** | **5116** | **5** | **530** |
| 18 | **AUR62029057** | **CqNAC17** | **NAC017: NAC domain-containing protein 17 \*** | **cq2_A** | **37120934** | **37126563** | **5630** | **4** | **524** |
| 19 | AUR62035514 | CqNAC21 | NAC021: NAC domain-containing protein 21/22 | cq5_B | 51399005 | 51402623 | 3619 | 2 | 153 |
| 20 | AUR62043497 | CqNAC21 | NAC021: NAC domain-containing protein 21/22 | cq12_A | 29606703 | 29610871 | 4169 | 2 | 155 |
| 21 | AUR62011853 | CqNAC21 | NAC021: NAC domain-containing protein 21/22 | cq8_A | 1858490 | 1862276 | 3787 | 2 | 145 |
| 22 | AUR62024698 | CqNAC21 | NAC021: NAC domain-containing protein 21/22 | cq9_B | 11061255 | 11065302 | 4048 | 3 | 149 |
| 23 | AUR62001365 | CqNAC25 | NAC025: NAC transcription factor 25 | cq7_A | 99573268 | 99576243 | 2976 | 3 | 307 |
| 24 | AUR62014768 | CqNAC25 | NAC025: NAC transcription factor 25 | cq1_B | 30397116 | 30398602 | 1487 | 3 | 343 |
| 25 | AUR62016954 | CqNAC25 | NAC025: NAC transcription factor 25 | cq2_A | 20121444 | 20123003 | 1560 | 3 | 327 |
| 26 | AUR62005821 | CqNAC29 | NAC029: NAC transcription factor 29 | cq14_A | 57750521 | 57756348 | 5828 | 5 | 288 |
| 27 | AUR62032816 | CqNAC29 | NAC029: NAC transcription factor 29 | cq6_B | 5793401 | 5794833 | 1433 | 3 | 272 |
| 28 | AUR62022690 | CqNAC30 | NAC030: NAC domain-containing protein 30 | cq10_A | 5723986 | 5732908 | 8923 | 5 | 571 |
| 29 | AUR62004648 | CqNAC30 | NAC030: NAC domain-containing protein 30 | cq1_B | 121459828 | 121461574 | 1747 | 3 | 334 |
| 30 | AUR62024101 | CqNAC31 | NAC031: Protein CUP-SHAPED COTYLEDON 3 | cq10_A | 59698378 | 59699514 | 1137 | 4 | 248 |
| 31 | AUR62035165 | CqNAC31 | NAC031: Protein CUP-SHAPED COTYLEDON 3 | cq3_B | 1657868 | 1658173 | 306 | 1 | 101 |
| 32 | AUR62043713 | CqNAC37 | NAC037: NAC domain-containing protein 37 | cq0_ | 14884377 | 14886152 | 1776 | 3 | 265 |
| 33 | AUR62043235 | CqNAC37 | NAC037: NAC domain-containing protein 37 | cq9_B | 41526 | 43301 | 1776 | 3 | 265 |
| 34 | AUR62034813 | CqNAC41 | NAC041: NAC domain-containing protein 41 | cq16_B | 11304423 | 11306000 | 1578 | 3 | 228 |
| 35 | AUR62029543 | CqNAC41 | NAC041: NAC domain-containing protein 41 | cq8_A | 26693056 | 26699989 | 6934 | 5 | 365 |
| 36 | AUR62026904 | CqNAC42 | NAC042: NAC domain-containing protein 42 | cq4_A | 44976454 | 44984863 | 8410 | 3 | 344 |
| 37 | AUR62040889 | CqNAC42 | NAC042: NAC domain-containing protein 42 | cq1_B | 72702858 | 72708102 | 5245 | 3 | 213 |
| 38 | AUR62004554 | CqNAC43 | NAC043: NAC domain-containing protein 43 | cq1_B | 120299216 | 120301237 | 2022 | 3 | 334 |
| 39 | AUR62019936 | CqNAC43 | NAC043: NAC domain-containing protein 43 | cq16_B | 72986058 | 72987726 | 1669 | 3 | 341 |
| 40 | AUR62022669 | CqNAC43 | NAC043: NAC domain-containing protein 43 | cq10_A | 5477634 | 5479368 | 1735 | 3 | 250 |
| 41 | AUR62000492 | CqNAC45 | NAC045: NAC domain-containing protein 45 | cq12_A | 5500539 | 5501585 | 1047 | 2 | 327 |
| 42 | AUR62004778 | CqNAC45 | NAC045: NAC domain-containing protein 45 | cq12_A | 25145721 | 25148970 | 4056 | 6 | 576 |
| 43 | AUR62003400 | CqNAC45 | NAC045: NAC domain-containing protein 45 | cq9_B | 1759783 | 1763912 | 4130 | 6 | 573 |
| 44 | AUR62037478 | CqNAC45 | NAC045: NAC domain-containing protein 45 | cq5_B | 62910115 | 62914170 | 3250 | 3 | 241 |
| 45 | AUR62015629 | CqNAC56 | NAC056: NAC transcription factor 56 | cq5_B | 3821692 | 3822921 | 1230 | 3 | 302 |
| 46 | AUR62029331 | CqNAC56 | NAC056: NAC transcription factor 56 | cq12_A | 51490439 | 51492813 | 2375 | 4 | 327 |
| 47 | **AUR62001531** | **CqNAC60** | **NAC60: NAC domain-containing protein 60 \*** | **cq7_A** | **101732643** | **101734168** | **1526** | **3** | **330** |
| 48 | AUR62007071 | CqNAC66 | NAC066: NAC domain-containing protein 66 | cq13_A | 5441599 | 5443027 | 1429 | 3 | 279 |
| 49 | AUR62016200 | CqNAC67 | NAC67: NAC domain-containing protein 67 | cq7_A | 75912843 | 75914874 | 2032 | 3 | 257 |
| 50 | AUR62036626 | CqNAC68 | NAC68: NAC domain-containing protein 68 | cq3_B | 56560424 | 56570523 | 10100 | 4 | 279 |
| 51 | AUR62025580 | CqNAC68 | NAC68: NAC domain-containing protein 68 | cq18_B | 24671246 | 24673006 | 1761 | 3 | 342 |
| 52 | AUR62025636 | CqNAC68 | NAC68: NAC domain-containing protein 68 | cq18_B | 23977517 | 23979442 | 1926 | 3 | 354 |
| 53 | AUR62015630 | CqNAC72 | NAC072: NAC domain-containing protein 72 | cq5_B | 3807048 | 3808199 | 1152 | 4 | 278 |
| 54 | AUR62022486 | CqNAC72 | NAC072: NAC domain-containing protein 72 | cq1_B | 96255823 | 96265494 | 9672 | 6 | 386 |
| 55 | AUR62029344 | CqNAC72 | NAC072: NAC domain-containing protein 72 | cq12_A | 51286101 | 51287415 | 1315 | 4 | 298 |
| 56 | AUR62021364 | CqNAC73 | NAC073: NAC domain-containing protein 73 | cq1_B | 16997945 | 17004953 | 7009 | 6 | 167 |
| 57 | AUR62033276 | CqNAC73 | NAC073: NAC domain-containing protein 73 | cq2_A | 52375299 | 52382203 | 6905 | 5 | 319 |
| 58 | AUR62031086 | CqNAC73 | NAC073: NAC domain-containing protein 73 | cq2_A | 5673254 | 5676404 | 3151 | 2 | 251 |
| 59 | AUR62014879 | CqNAC73 | NAC073: NAC domain-containing protein 73 | cq1_B | 32318489 | 32327115 | 8627 | 2 | 252 |
| 60 | AUR62026259 | CqNAC73 | NAC073: NAC domain-containing protein 73 | cq14_A | 24108385 | 24115692 | 7308 | 4 | 191 |
| 61 | AUR62043434 | CqNAC73 | NAC073: NAC domain-containing protein 73 | cq0_ | 19574734 | 19575424 | 691 | 3 | 138 |
| 62 | AUR62036063 | CqNAC73 | NAC073: NAC domain-containing protein 73 | cq15_A | 45801852 | 45808337 | 6486 | 3 | 204 |
| 63 | **AUR62041525** | **CqNAC78** | **NAC078: NAC domain-containing protein 78 \*** | **cq5_B** | **57361157** | **57364321** | **3165** | **6** | **550** |
| 64 | AUR62043491 | CqNAC78 | NAC078: NAC domain-containing protein 78 | cq12_A | 28893343 | 28901372 | 8030 | 4 | 393 |
| 65 | **AUR62041539** | **CqNAC78** | **NAC078: NAC domain-containing protein 78 \*** | **cq12_A** | **20501797** | **20505411** | **3615** | **6** | **556** |
| 66 | AUR62043741 | CqNAC78 | NAC078: NAC domain-containing protein 78 | cq1_B | 84648679 | 84662099 | 13421 | 4 | 355 |
| 67 | AUR62006885 | CqNAC81 | NAC081: Protein ATAF2 | cq5_B | 74997825 | 75001903 | 4079 | 4 | 329 |
| 68 | AUR62006407 | CqNAC82 | NAC082: NAC domain-containing protein 82 | cq7_A | 90788443 | 90789561 | 1119 | 1 | 372 |
| 69 | AUR62029755 | CqNAC82 | NAC082: NAC domain-containing protein 82 | cq17_B | 59895805 | 59896911 | 1107 | 1 | 368 |
| 70 | AUR62006837 | CqNAC82 | NAC082: NAC domain-containing protein 82 | cq5_B | 75576113 | 75577156 | 1044 | 2 | 319 |
| 71 | AUR62000493 | CqNAC82 | NAC082: NAC domain-containing protein 82 | cq12_A | 5522447 | 5523436 | 990 | 1 | 329 |
| 72 | AUR62028872 | CqNAC83 | NAC083: NAC domain-containing protein 83 | cq6_B | 3320845 | 3322156 | 1312 | 4 | 257 |
| 73 | AUR62005686 | CqNAC83 | NAC083: NAC domain-containing protein 83 | cq14_A | 55484398 | 55485672 | 1275 | 4 | 256 |
| 74 | AUR62018752 | CqNAC86 | NAC086: NAC domain-containing protein 86 | cq16_B | 75482814 | 75484845 | 2032 | 4 | 325 |
| 75 | AUR62007247 | CqNAC86 | NAC086: NAC domain-containing protein 86 | cq13_A | 3165461 | 3167486 | 2026 | 4 | 327 |
| 76 | AUR62010080 | CqNAC86 | NAC086: NAC domain-containing protein 86 | cq15_A | 5122099 | 5126510 | 4412 | 4 | 352 |
| 77 | AUR62013074 | CqNAC86 | NAC086: NAC domain-containing protein 86 | cq17_B | 79021049 | 79025517 | 4469 | 4 | 352 |
| 78 | **AUR62020251** | **CqNAC89** | **NAC089: NAC domain-containing protein 89 \*** | **cq18_B** | **32521438** | **32523891** | **2454** | **5** | **430** |
| 79 | AUR62001522 | CqNAC90 | NAC090: NAC domain-containing protein 90 | cq7_A | 101656423 | 101660122 | 3700 | 4 | 268 |
| 80 | AUR62020261 | CqNAC90 | NAC090: NAC domain-containing protein 90 | cq18_B | 32661219 | 32663097 | 1879 | 3 | 258 |
| 81 | AUR62006223 | CqNAC90 | NAC090: NAC domain-containing protein 90 | cq15_A | 2810155 | 2814517 | 4363 | 3 | 276 |
| 82 | AUR62036991 | CqNAC90 | NAC090: NAC domain-containing protein 90 | cq17_B | 56071227 | 56073064 | 1838 | 2 | 151 |
| 83 | AUR62039815 | CqNAC94 | ANAC094: Putative NAC domain-containing protein 94 | cq4_A | 38381888 | 38383418 | 1531 | 3 | 385 |
| 84 | AUR62034400 | CqNAC94 | ANAC094: Putative NAC domain-containing protein 94 | cq1_B | 55324061 | 55325551 | 1491 | 3 | 385 |
| 85 | AUR62027152 | CqNAC94 | ANAC094: Putative NAC domain-containing protein 94 | cq15_A | 1809369 | 1810776 | 1408 | 3 | 222 |
| 86 | AUR62012948 | CqNAC94 | ANAC094: Putative NAC domain-containing protein 94 | cq17_B | 80277875 | 80278498 | 624 | 2 | 146 |
| 87 | AUR62040120 | CqNAC94 | ANAC094: Putative NAC domain-containing protein 94 | cq6_B | 34103882 | 34105819 | 1938 | 5 | 390 |
| 88 | **AUR62032488** | **CqNAC94** | **ANAC094: Putative NAC domain-containing protein 94 \*** | **cq14_A** | **41611790** | **41618469** | **6680** | **8** | **516** |
| 89 | AUR62038748 | CqNAC98 | NAC098: Protein CUP-SHAPED COTYLEDON 2 | cq1_B | 106129053 | 106135097 | 6045 | 4 | 294 |
| 90 | AUR62020634 | CqNAC98 | NAC098: Protein CUP-SHAPED COTYLEDON 2 | cq4_A | 16178718 | 16185086 | 6369 | 4 | 297 |
| 91 | AUR62019477 | CqNAC98 | NAC098: Protein CUP-SHAPED COTYLEDON 2 | cq12_A | 48919545 | 48921791 | 2247 | 3 | 400 |
| 92 | AUR62036158 | CqNAC98 | NAC098: Protein CUP-SHAPED COTYLEDON 2 | cq5_B | 11161825 | 11164198 | 2374 | 5 | 324 |
| 93 | AUR62034435 | CqNAC100 | NAC100: NAC domain-containing protein 100 | cq1_B | 56919532 | 56920931 | 1400 | 3 | 341 |
| 94 | AUR62018102 | CqNAC100 | NAC100: NAC domain-containing protein 100 | cq4_A | 35692016 | 35693362 | 1347 | 3 | 333 |
| 95 | AUR62021652 | CqNAC100 | NAC100: NAC domain-containing protein 100 | cq8_A | 9672471 | 9674125 | 1655 | 3 | 361 |
| 96 | AUR62008438 | CqNAC100 | NAC100: NAC domain-containing protein 100 | cq16_B | 4681091 | 4683124 | 2034 | 3 | 364 |
| 97 | AUR62021621 | CqNAC100 | NAC100: NAC domain-containing protein 100 | cq8_A | 10126416 | 10133113 | 6698 | 4 | 198 |
| 98 | **AUR62028103** | **CqNTL9** | **NTL9: Protein NTM1-like 9 \*** | **cq11_B** | **68788278** | **68792879** | **4602** | **6** | **630** |
| 99 | **AUR62005966** | **CqNTL9** | **NTL9: Protein NTM1-like 9 \*** | **cq7_A** | **70111368** | **70115912** | **4545** | **6** | **631** |
| 100 | AUR62013541 | CqSMB | SMB: Protein SOMBRERO | cq3_B | 65117611 | 65120613 | 3003 | 3 | 346 |
| 101 | AUR62026741 | CqSMB | SMB: Protein SOMBRERO | cq10_A | 20749557 | 20757214 | 7658 | 4 | 283 |
| 102 | AUR62005772 | CqFEZ | FEZ: Protein FEZ | cq14_A | 56770446 | 56773311 | 2866 | 3 | 446 |
| 103 | AUR62044373 | CqFEZ | FEZ: Protein FEZ | cq6_B | 5175069 | 5177844 | 2776 | 3 | 454 |
| 104 | AUR62017681 | CqBRN2 | BRN2: Protein BEARSKIN2 | cq0_ | 137754423 | 137760607 | 6185 | 4 | 418 |
| 105 | AUR62000898 | CqNAM-B1 | NAM-B1: NAC transcription factor NAM-B1 | cq12_A | 10832642 | 10835485 | 2844 | 3 | 418 |
| 106 | AUR62006836 | CqNAC103 | NAC103: NAC domain containing protein 103 | cq5_B | 75580825 | 75581836 | 1011 | 2 | 315 |
| 107 | AUR62004773 | CqNAC28 | NAC domain containing protein 28 | cq5_B | 62725181 | 62726259 | 1078 | 2 | 140 |

Quinoa NAC genes encode proteins ranging in size from 101 to 631 amino acids (aa) in length, with an average of 322 aa.

Several NAC TFs are present as membrane-bound transcription factors (MTFs) (15). To identify membrane associated NAC TFs in quinoa, we scanned all of the full length CqNAC peptide sequences for the presence of α-helical transmembrane (TM) motives using the TMHMM server v.2.0. (http://www.cbs.dtu.dk/services/TMHMM/). We identified nine putative membrane-bound CqNACs proteins containing α-helical TM motives in the C-terminal region (Table 1).

The number of CqNACs we identified in this study is less than the number of NACs in Arabidopsis and rice. We expected at least double the number of NACs due to the tetraploid nature of the quinoa genome. However, we found that the total number of predicted TFs in the genome of quinoa is in fact even slightly less than the total number of TFs in Arabidopsis (2093 CqTFs in quinoa compared with 2296 TFs in Arabidopsis, S1 Fig, S1 Table). The number of TFs in each family is almost 0.8-fold (median) compared with Arabidopsis, with only two families showing 3 and 5 times more TFs in quinoa compared with Arabidopsis i.e., the FAR1 and LFY families, respectively. Moreover, fewer sequences belonging to the heat shock proteins family 70 (HSP70) were identified in quinoa compared with Arabidopsis (quinoa has 16 HSP70 members, while Arabidopsis has 18 members) (26). However, by comparing the number of CqNAC genes in quinoa with its close relatives, quinoa has more NACs than spinach (*Spinacia oleracea*) and sugar beet (*Beta vulgaris*), which have 80 and 59 NACs genes, respectively.

### 2.2 Chromosomal location and gene duplication of quinoa *CqNAC* genes

In total, 103 CqNACs are localized to the 18 chromosomes of quinoa and only four CqNACs could not be mapped to any chromosome and are therefore assigned to chromosome zero (Table 1). CqNACs genes appear to be equally distributed between sub-genome A and B (S2a Fig); however, CqNACs are unequally distributed across the chromosomes (S2b Fig). The largest number of CqNACs genes are localized to chromosome 1 (13 CqNACs, ∼12.38 %), followed by chromosome 12 (12 CqNACs, ∼11.43 %), while chromosome 11 and chromosome 13 have the smallest number of CqNACs (only two CqNACs ∼1.9 % are localized to chromosome 11 and three CqNACs ∼2.86 % are localized to chromosome 13) (S2b Fig).

We noticed that in most cases, we found two or more quinoa *CqNAC* genes for every Arabidopsis orthologue and thus we grouped quinoa CqNACs as duplicated and un-duplicated genes. The duplicated group consists of 30 duplicated CqNAC genes (each gene has from 2 to 7 copies) giving a total number of 97 genes distributed between subgenome A and subgenome B. The un-duplicated CqNACs consist of 10 CqNACs genes (Fig 2a,b). In total, only 40 Arabidopsis NAC genes have orthologues in the genome of quinoa (36.5% of Arabidopsis NACs).

**Fig 2.**
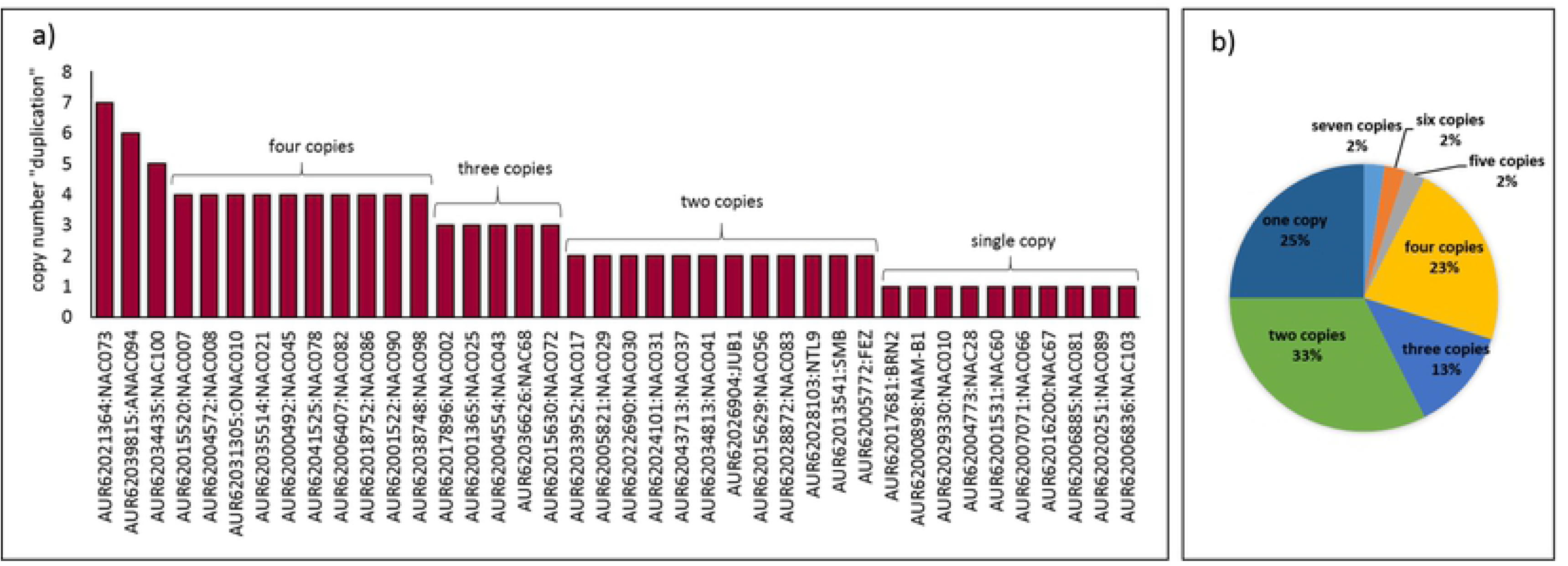
Duplicated and non-duplicated quinoa *CqNAC* genes. a) Copy number of each quinoa *CqNAC* gene. b) Pie chart shows the percentage of duplicated and unduplicated *CqNACs*. The percentage of genes are calculated based on the total number of *CqNACs* orthologues to Arabidopsis *NACs* (40 *NAC* genes).

We performed synteny analysis using MCScanX to reveal the relationship between the duplicated CqNACs, and to determine if these duplicated genes form homologous pairs and if these homologs are located between subgenomes A and B (to form homoeologous pairs). Synteny analysis revealed that different types of homology occurs between the duplicated genes, which we classify as: (1) genes that form a homoeologous pair located in the same phylogenetic cluster (S3a Table) – in quinoa, this applies to 52 genes (26 pairs, 48.6% of all of the genes); (2) genes that appear to form a homoeologous pair but are located in different phylogenetic clusters - 18 genes in quinoa (16.8%), (S3b Table); (3) genes that have more than one homolog located in different clusters - 10 genes in quinoa (9.3%) (S3c Table); and (4) genes that have no homolog according to our analyses - 27 genes in quinoa (25.2%) (S3d Table). Although some of these genes are duplicated, such as the orthologue of *AtJUB1*, none of the duplicated copies have a homoeolog. The absence of some homoeolog copies in some of the *CqNAC* genes suggests that gene loss occurred during the evolution of quinoa, causing some gene loss in the NAC family. To confirm this, the sequences of the two progenitor genomes needs to be investigated. Similar findings have been observed in the WRKY TF family in quinoa, where some of the homologous copies of some of the WRKY genes have been lost (25).

### 2.3 Phylogenetic analysis of the NAC TFs family in quinoa and Arabidopsis

We constructed a Neighbor-Joining (N-J) phylogenetic tree to study the relationship between quinoa CqNACs. The NAC domain of 107 quinoa NAC proteins was used in the phylogenetic construction. From the phylogenetic tree (and according to the bootstrap values), quinoa NAC proteins can be classified into two major groups (Fig 3a). Group I has the largest number of CqNACs (96 CqNACs) and is subdivided into 15 subgroups (Fig 3b). Group II contains 11 CqNACs.

**Fig 3.**
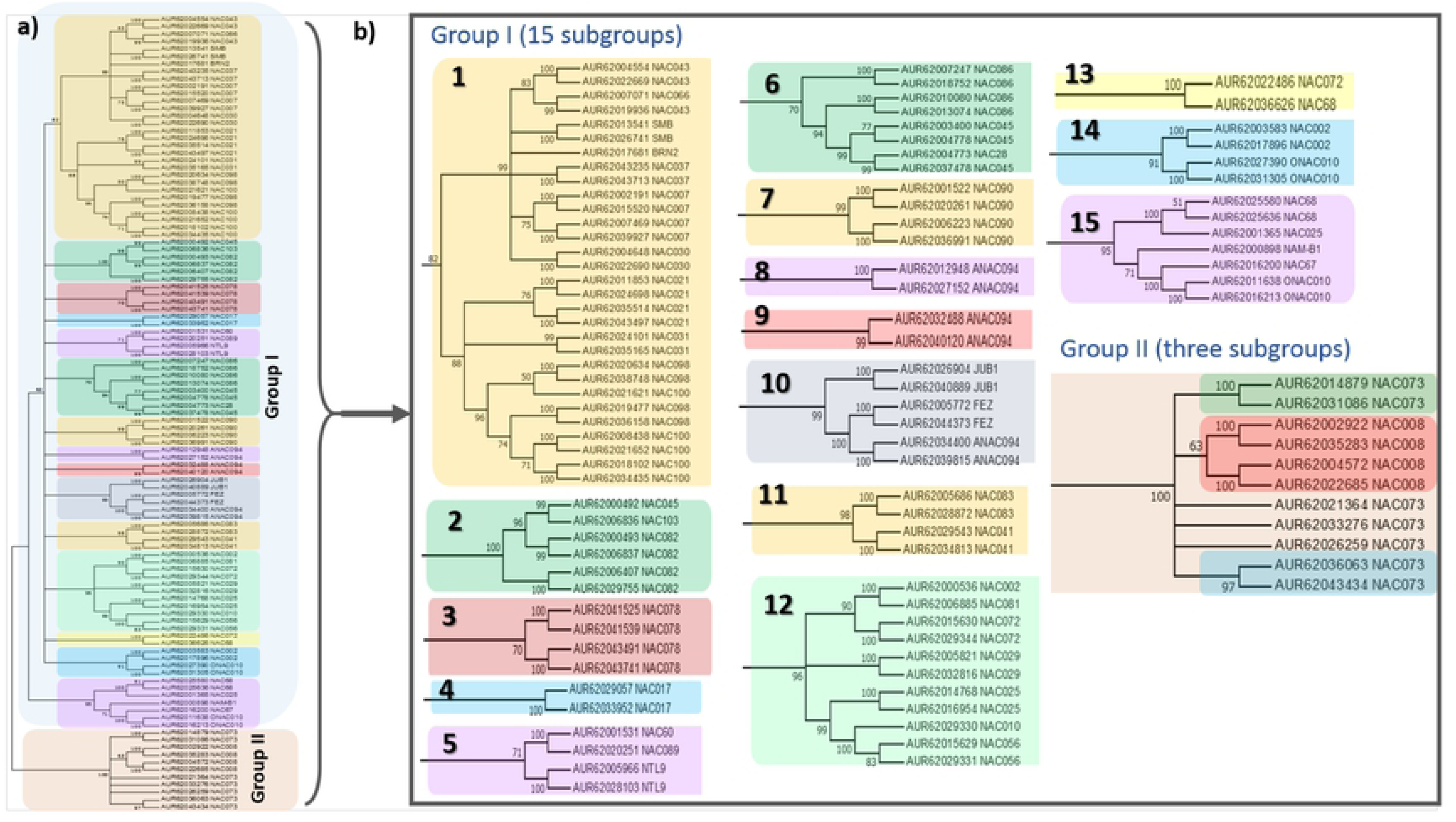
Phylogenetic tree of quinoa CqNACs. a) The phylogenetic tree was constructed in MEGA 7.0 using the Neighbour-Joining (NJ) method with 1000 bootstrap interactions based on the multiple sequence alignment of 107 amino acid sequences of NAC domains belonging to the NAC proteins from *Chenopodium quinoa*. Nodes that have a bootstrap value of less than 50 are collapsed. The tree divided CqNACs into two major groups: Group I (with blue background) and Group II (with pink background). Group I is further subdivided into 15 subgroups that have different background colours. b) Details of the subgroups from the phylogenetic tree in (a).

Generally, most of the quinoa CqNACs form sister pairs (there are 49 pairs) except for nine CqNACs that occur as a single CqNAC (Fig 3b). These results indicate that genes in each subgroup might originate from the same duplication event. Similar phylogenetic topologies have been also found in other plants with duplicated genomes, such as in switchgrass (28). For most NACs, closely related members in the same phylogenetic subgroup share a similar exon/intron structure (number and length), with few exceptions (Fig 4).

**Fig 4.**
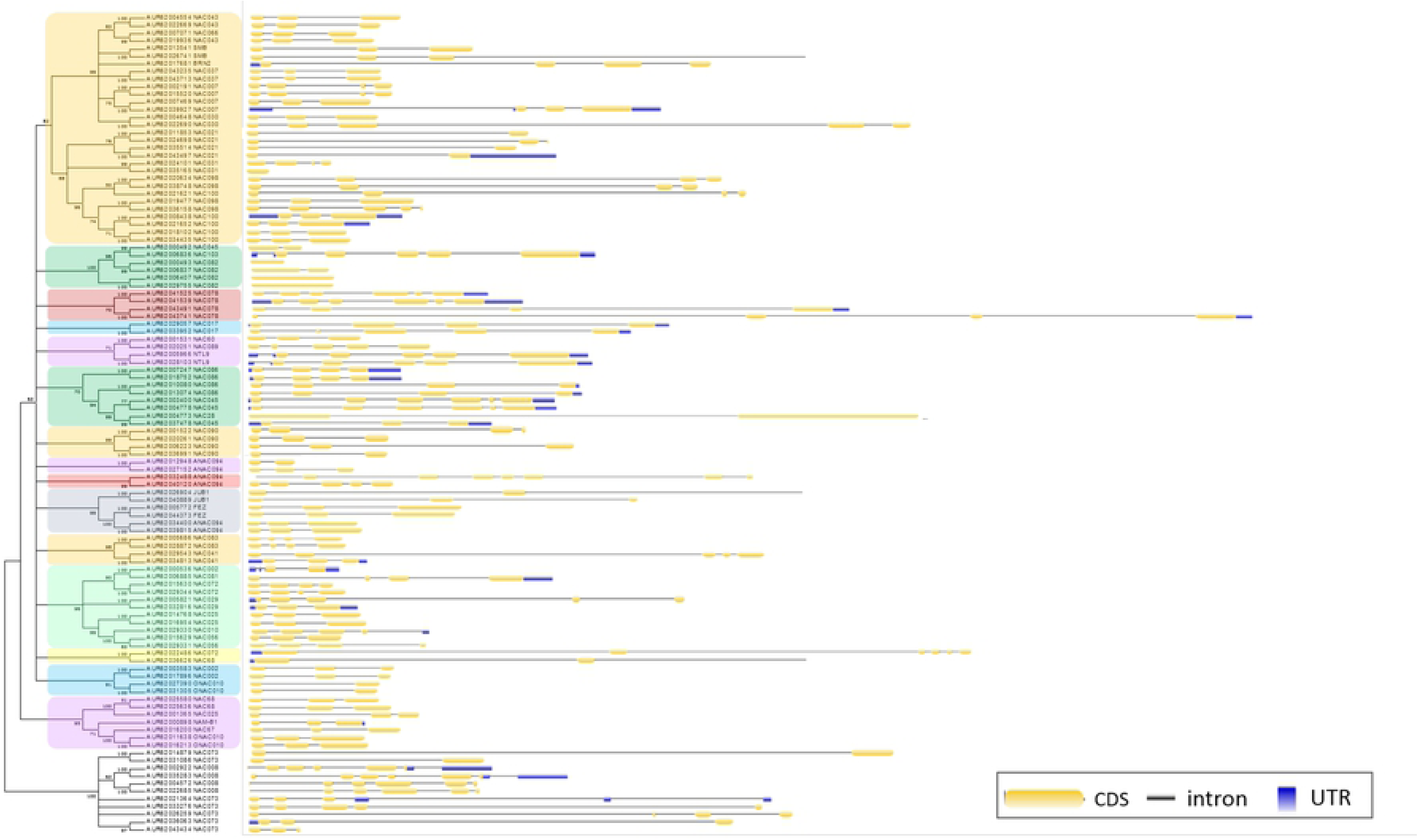
Exon/ intron structure of quinoa *CqNAC* genes belonging to different phylogenetic subgroups.

To find the phylogenetic relationship between quinoa and Arabidopsis, we constructed another phylogenetic tree, in this case from the alignments of the full-length sequences of NAC proteins from Arabidopsis (110 seq.) and quinoa (107 seq.). (Fig 5). The phylogenetic tree divided NACs into different sub-groups, consistent with the previous classification (34). The bootstrap values were sometimes low; however, this was also reported in previous studies (35–39). NAC TFs from the same phylogenetic group are likely to have a similar function. For instance, the NAM subfamily includes NACs that function in shoot boundary formation (CUC1/2/3) (40, 41), while NACs belonging to subfamily OsNAC7 are involved in secondary wall formation (Vascular related NAC-Domain proteins “VNDs” (42) and in root development (SMBs (43)). The OsNAC8 family are another example, which includes NACs involved in stress response (ANAC019, ANAC055 and ANAC072) (44). Interestingly, there is a subgroup of Arabidopsis NACs that are absent in quinoa (subgroups 8, 16 and 18), might have been lost during quinoa evolution. Subgroups 8, 16 and 18 contain mostly unknown NACs, where the function has not been described yet. There are only four genes previously described, ANAC005 involved in xylem formation (45), NTM1 involved in cell division (46) and two genes involved in abiotic stress, NTM2 and NAC67 (47, 48). Quinoa has a high tolerance to salinity and water deficit; it is possible, that the absence of these genes contributes to quinoa’s tolerance. There are only two subgroups of NACs for which quinoa has a larger number of members than Arabidopsis, and there are three groups that quinoa is missing (S4 Table). This may explain why quinoa has less NACs than Arabidopsis.

**Fig 5.**
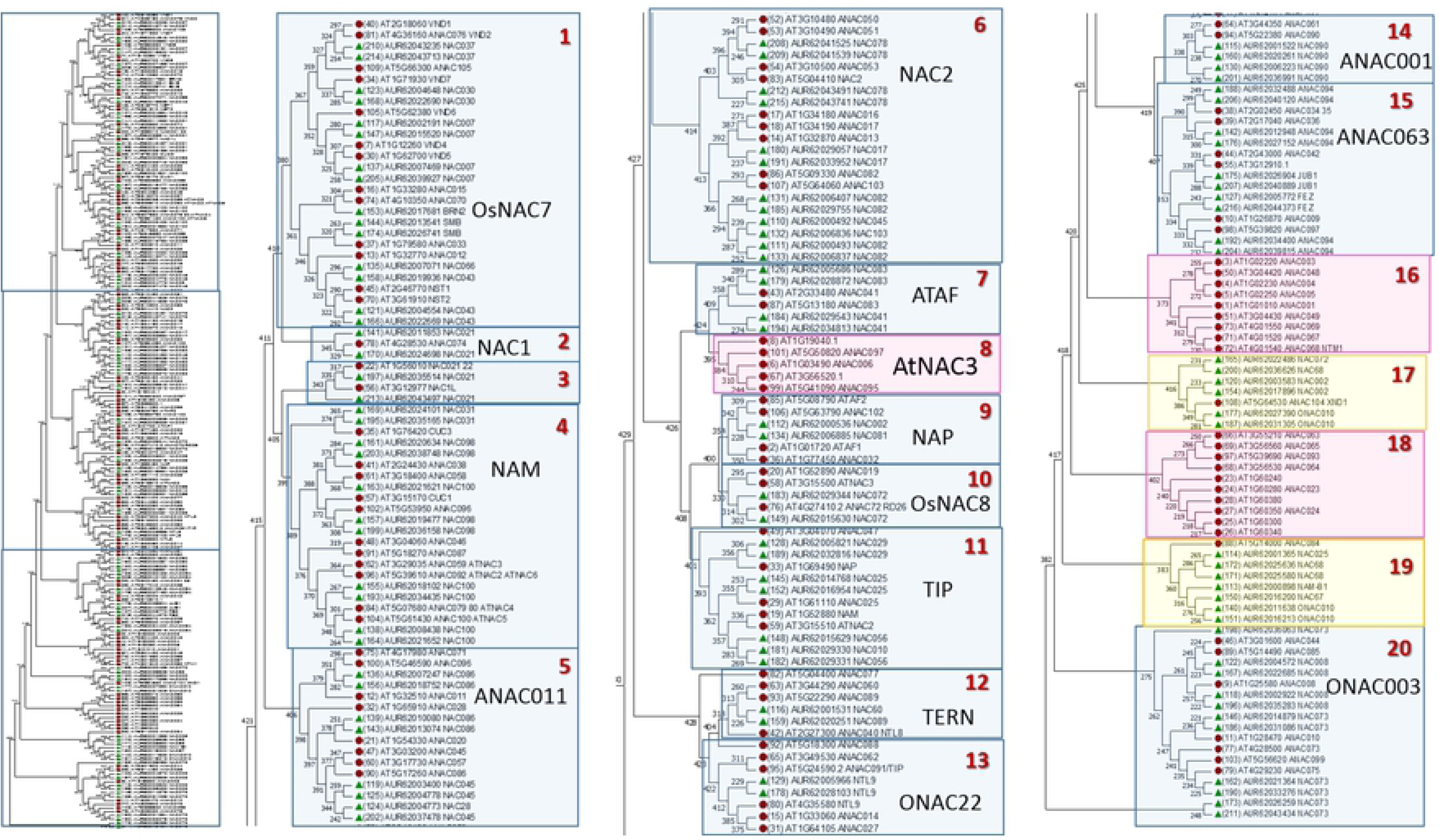
Phylogenetic relationship of NAC proteins from Arabidopsis and Quinoa. The phylogenetic tree was constructed in MEGA 7.0 using the Neighbor-Joining (NJ) method with 1000 bootstrap interactions based on multiple sequence alignment of 107 amino acid sequences of full-length peptide sequence from *Chenopodium quinoa* and 110 sequences from Arabidopsis thaliana. The tree was divided into different subgroups and named according to the classification of OoKa et al. (2013). Red circles represent NAC peptides from Arabidopsis and green triangles represent peptides from quinoa. Blue boxes represent subgroups that have members from both Arabidopsis and quinoa, yellow boxes represents subgroups that have larger number of quinoa NACs than Arabidopsis NACs and pink boxes represents subgroups that have members from Arabidopsis only.

### 2.4 Transcriptional analyses of quinoa *CqNAC* genes

#### 2.4.1 Expression of duplicated genes (functionalization of duplicated genes)

The high proportion of gene duplication among quinoa *CqNAC* genes raises the question of their functional redundancy. During the evolutionary process, duplicated genes may have obtained non-functionalization, neo-functionalization or sub-functional redundancy (49), which could be indicated by differences in their patterns of expression. Thus, we investigated the expression of duplicated *CqNAC*s in the shoots and roots of quinoa plants grown in soil and in a hydroponic system under normal growth conditions. We used the available RNAseq data for quinoa accession QQ74 assessed under different stress conditions, such as heat, drought, salinity and low-phosphate. First, to determine if all of the *CqNAC*s that we identified in this study were expressed, we calculated their cumulative expression in all of the samples and all of the conditions. We found that five genes were not expressed at all under any of the conditions used in this work, and four genes were expressed only at very low levels (Fragments Per Kilobase of transcript per Million mapped reads, FPKM<1) (S5 Table).

Expression analysis of the duplicated genes showed a different expression pattern between the shoots and the roots among the duplicated copies (Fig 6). Some of the duplicated *CqNAC* genes have a similar expression pattern, which could suggest a similar or redundant function. Other duplicated *CqNAC* genes have different expression patterns, suggesting that some of the duplicated copies may have developed different functions. Similar expression patterns were also observed in diploid cotton *Gossypium raimondii* (30) and Tartary buckwheat (*Fagopyrum tataricum*) (50). We also noticed that not all of the duplicated copies are expressed. For example, in two of the genes that are triplicated and one of the genes that are found in four copies, only two copies are expressed and the other two copies are not expressed.

**Fig 6.**
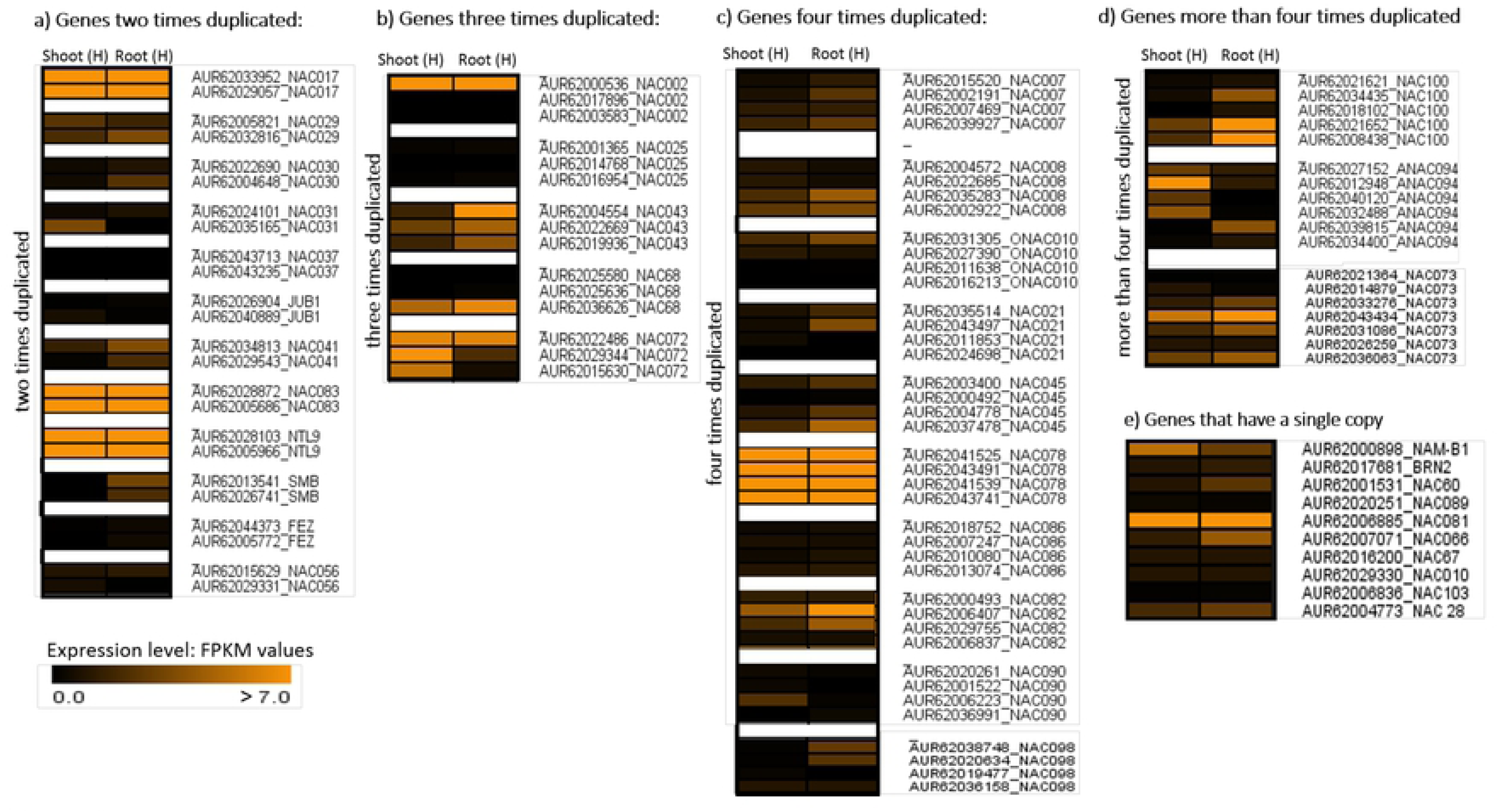
Heat map of the expression of duplicated *CqNAC* in the shoots and roots of quinoa. Quinoa plants were grown in hydroponic system under normal growth conditions. a) Genes duplicated two times, b) Genes duplicated three times, c) Genes duplicated four times, d) Genes duplicated more than four times, and e) Genes that have a single copy. The corresponding expression values are in S6 Table.

NAC TFs have long been known as an important TF family that are involved in regulating plant growth, development and responses to stress (1, 2, 51–53). To detect NACs that are potentially involved in root and shoot development, we identified the expression of all of the 107 putative quinoa NAC TFs genes in the shoots and roots of quinoa plants grown under growth conditions in soil as well as in hydroponic conditions (S6 Table). To find the genes that are potentially involved in root or shoot development, the expression of *CqNAC*s in the shoots was compared to their expression in the roots. We chose genes that were significantly differentially expressed (DE) between shoot and root (FDR=0.05), as these might be involved in regulating root or shoot development. Those *CqNACs* comprise 24 genes (19 genes being more expressed in roots “root specific” and five genes being more induced in the shoots “shoot specific”) (Fig 7 and S3 Fig and S6 Table). The expression of *CqNAC*s in the shoots and roots of plants grown in soil and in hydroponic showed a consistent expression pattern. However, there are a few differences in the number of significantly differentially expressed (DE) genes. Among those NAC genes that are significantly more expressed in the roots than the shoots (in both soil and hydroponic) is the *AUR62013541* gene, which is the ortholog of the well-characterized *SOMBRERO* (*SMB*) gene. *SMB* has been shown to be involved in controlling the reorientation and timing of cell division in root stem cells (43). *AUR62000536* is a second example, the orthologue of the Arabidopsis *AtNAC2* gene. *AtNAC2* is specifically expressed in the roots and significantly increases the number of lateral root formation in response to ethylene and auxin signalling in Arabidopsis (54). In addition, *AUR62032816* (the orthologue of *ANAC02*9) is strongly and significantly induced in the roots compared to the shoots. *ANAC029* was found to be involved in root morphogenesis in *Medicago truncatula* (55). *AUR6200455*4 is another example, which is the orthologue of Arabidopsis *ANAC043*. In Arabidopsis, *ANAC043* belongs to secondary wall NAC genes (SWN), which are also known as *secondary wall thickening promoting factor1* (*NST1*); however, its role in root growth and development has not yet been identified (56). With respect to shoot related *CqNACs*, we identified only five *CqNACs* to be significantly more expressed in the shoots than the roots, i.e., the two orthologues of *ANAC072* (*AUR62015630*, *AUR62029344*), and the two orthologs of *ANA0C94* (*AUR62032488*, *AUR62040120*) and the orthologue of *NAM-B* (*AUR62000898*). The function of these genes in shoot formation has not been identified before. Thus, future studies should test if those genes are indeed involved in shoot formation or shoot related processes in quinoa.

**Fig 7.**
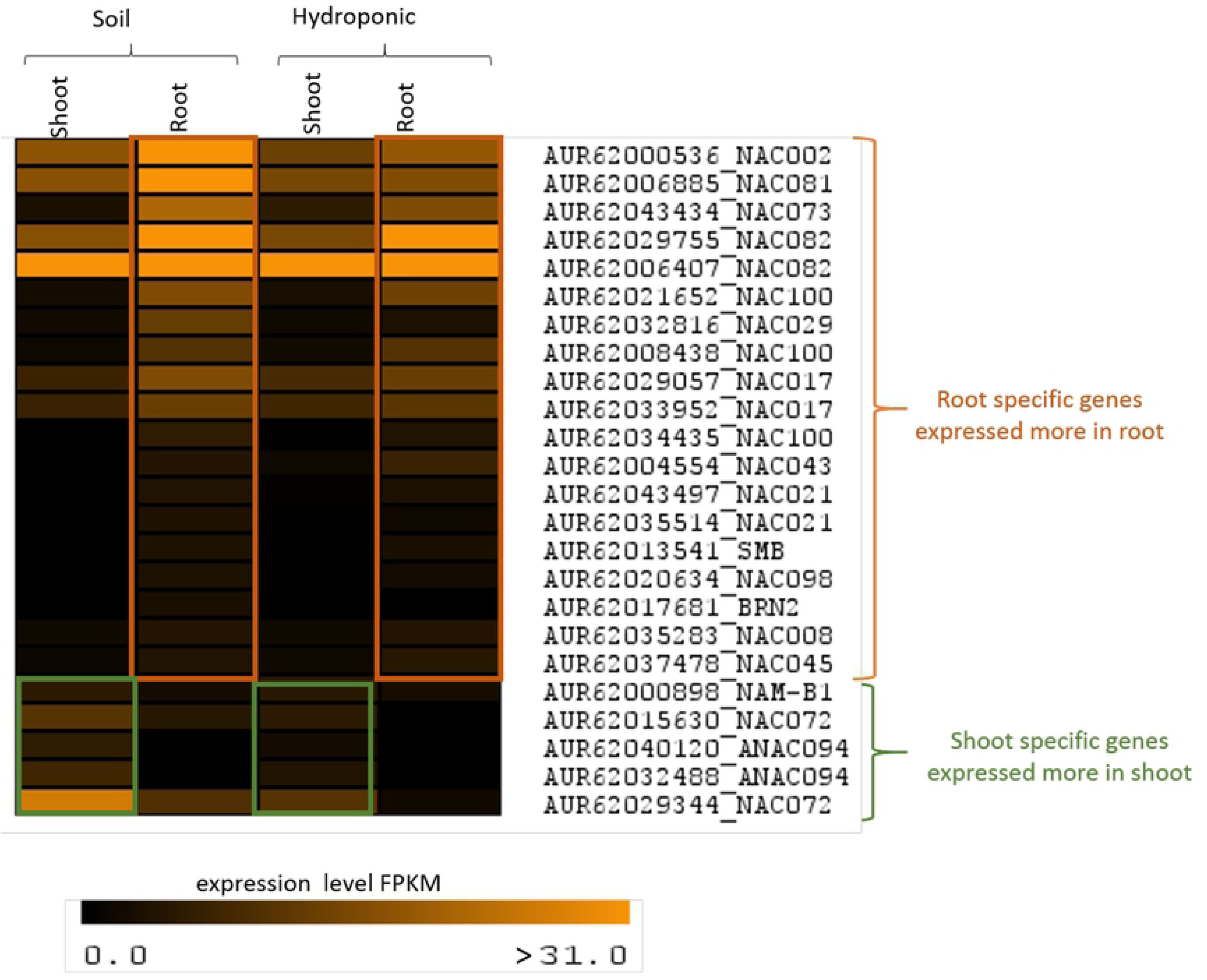
Heat map of the expression of *CqNAC* significantly differentially expressed (DE) between roots and shoots. Expression level (in FPKM) of significantly DE genes between shoot and root of plants grown in a hydroponic system or soil. Significant difference is calculated at FDR=0.05.

#### 2.4.2 Expression of quinoa *CqNAC* TFs in response to abiotic stresses

NAC TFs have been identified as an important regulator of stress responses, and given that quinoa has a high stress tolerance to abiotic stresses, we hypothesize that NAC TFs are involved in quinoa’s adaptation to stress tolerance. Here, we examine the expression of all of the putative quinoa NACs in response to different abiotic stresses (salt, drought, heat and phosphate starvation) using RNAseq previously generated (27). Around 35-48 *CqNAC*s are DE in the roots or shoots of quinoa plants exposed to one of the abiotic stresses examined in this study. However, the largest number of genes being DE were in the roots of the plants that were exposed to phosphate starvation (Fig 8a).

**Fig 8.**
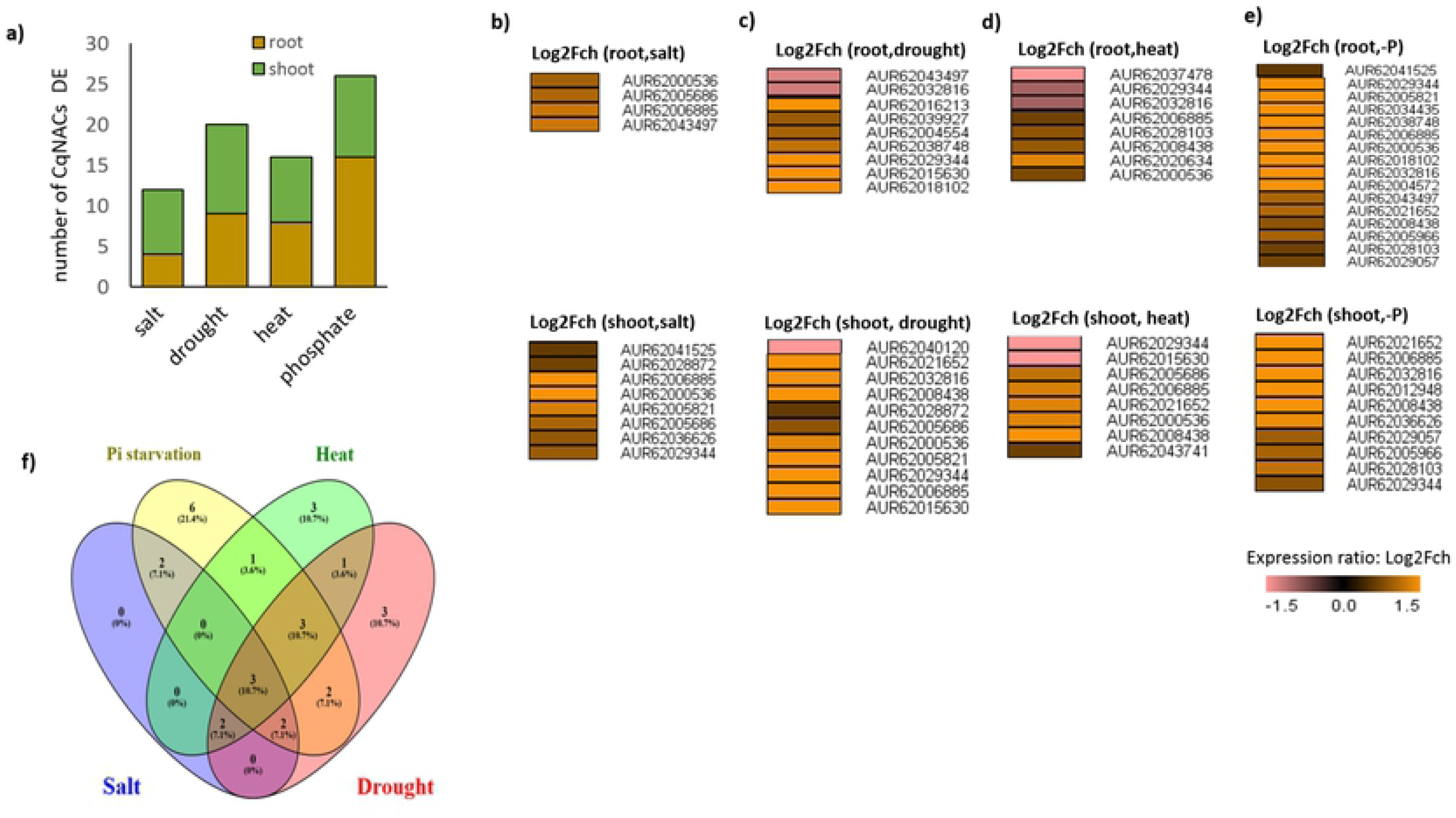
Quinoa NACs differentially expressed in response to different abiotic stresses. a) Number of genes differentially expressed in the shoots and roots of quinoa plants exposed to different abiotic stresses, including salt, drought, heat and phosphate starvation. b) Heat map of *CqNAC*s DE after salt stress. c) Heat map of *CqNAC*s DE after drought stress. d) Heat map of *CqNAC*s DE after heat stress. e) Heat map of *CqNAC*s DE after phosphate starvation. The significance between stress and control is calculated at FDR=0.05. f) Venn diagram illustrating the overlap between different stress-responsive *CqNAC* genes (only significant DE genes).

NAC genes are known to be involved in salt stress response and tolerance in Arabidopsis and many other plant species (57–60). Here, we identified nine genes in quinoa that were significantly upregulated in the shoots, roots or both in response to salt stress (5 genes in shoot, 1 gene in root and 3 genes in both shoot and root) (Fig 8b). Interestingly, some of these genes are orthologues of NACs genes, which have been previously identified as salt responsive genes and are known to play a role in salinity tolerance when they are overexpressed in plants. For example, *AUR62028872* and *AUR62005686* are two orthologues of the Arabidopsis *ANAC083* (*VND-INTERACTING 2* or *VIN2*), a negative regulator of xylem vessel formation. In Arabidopsis, *ANAC083* was induced by salt treatment and its overexpression enhanced salinity tolerance by directly binding to the promoters of two important genes involved in salt stress response (61). Our data showed a similar trend as the two quinoa orthologues of *ANAC083* were significantly induced by salt stress. *AtNAP* (*ANAC029)* was identified as another NAC TF gene that is induced by salt stress (62). *AtNAP* functions as a negative regulator of salt stress response via repressing the expression of *AREB1* and thus its overexpression increases the plants’ sensitivity to salt stress (62). We found a similar expression trend in quinoa as the quinoa orthologue of *AtNAP*, *AUR62005821*, was also induced by salt stress in both shoot and root. Two other genes that are also significantly upregulated by salt treatment are *AUR62000536,* the *NAC2* orthologue and *AUR62029344*, the A*NAC072* orthologue – those two genes were found to be induced by salt stress, and plants were found to have enhanced stress tolerance when they were overexpressed separately (10, 63). Another example is *AUR62006885,* the *ATAF2* orthologue, which has been previously identified to be induced by salt stress and pathogen attack (64, 65). Here we report a similar expression pattern in quinoa as it is also induced in the shoots and roots of quinoa plants in response to salt stress.

Some salt responsive *CqNAC* genes, such as *AUR62036626*, *AUR62041525* and *AUR62043497* (the orthologues of Arabidopsis *ANAC068*, *ANAC078* and *ANAC021*, respectively), are significantly DE in response to salt, but their roles in response to salt stress have not been studied before. These genes have the potential to be involved in salt stress response and adaptation in quinoa.

Quinoa is known to have a high drought tolerance and many NACs have been found to be involved in drought tolerance in rice and Arabidopsis (57–60). After drought treatment, expression of sixteen *CqNAC* genes were significantly different between the stress and control in the shoots or roots, or both (Fig 8c). Some of these significantly DE genes are orthologous to well-known drought responsive NAC genes in other plant species. For example, in Arabidopsis, overexpression of *RD26*/*ANAC072* and *ATAF1*/*ANAC002* confers drought tolerance (44, 66). We found that in quinoa the *ATAF-1* orthologue *AUR62000536* and the two orthologues of *ANAC072 AUR62015630* and *AUR62029344* were upregulated in response to drought stress, which is consistent with their role in drought tolerance as for Arabidopsis (44, 66). *AUR62005821* and *AUR62032816*, the two orthologues of Arabidopsis *AtNAP*, present two further examples, which we found to be significantly upregulated in the shoots and downregulated in the roots in response to drought stress. In rice, *OsNAP* functions as a positive regulator of drought tolerance (59). In addition, the wheat *TaNAP29* enhances drought tolerance in transgenic Arabidopsis (60). We identify ten genes that are significantly differentially expressed, but their roles in response to drought stress have not been previously identified.

The focus of the role of NAC TFs in heat response has only recently emerged and there are only a few reports focusing on their role in heat tolerance. Thus, their role has not been studied as thoroughly as other stresses (53, 67–70). Here, we identified 13 genes that were significantly differentially expressed in response to heat stress in the shoots or roots or both. Interestingly, *AUR62015630* and *AUR62029344* (the two orthologues of *ANAC072/RD26*) were downregulated in both the shoots and roots. In our previous study of the transcriptomic analysis of NAC TFs in response to heat priming in Arabidopsis, we found that *RD26* and *AtNAP* were responsive to heat priming (Alshareef *et al*, unpublished). This suggests the possible role of NAC TF genes in regulating heat stress response in quinoa.

Adequate supply of inorganic phosphate (P_i_) is important for plant growth and development, whereby low levels of P_i_ in soil adversely affect plant growth and development. Thus, plants have evolved several regulatory mechanisms to adapt to phosphorus deficiency and optimize P_i_ uptake and assimilation (57). Here, we found that the largest number of quinoa NAC TF genes was induced by phosphate deficiency treatment. Nineteen genes were significantly upregulated in the roots or shoots or both (16 *CqNAC* genes were upregulated in the roots and 10 *CqNACs* were upregulated in in the shoots in response to phosphate deficiency (Fig 8e). In Arabidopsis, it has been reported that five NAC genes are induced by more than a 2 fold change in response to P-starvation, namely *At3g15500/AtNAC3*, *At3g04070/ANAC047*, *At2g43000/JUB1*, *At1g52890/ANAC019*, *At1g02220/ANAC003* (71–73) and (Hammond and Bennet, (AT-O61 Genevestigator). However, none of these gene orthologues in quinoa are significantly differentially expressed in our data.

From the above results on the transcriptional responses of quinoa NACs to different abiotic stresses, we found there are three *CqNAC*s that are differentially expressed in response to all of the stresses, i.e., *AUR62029344*, *AUR62000536* and *AUR62006885*, which are the orthologues of *ANAC072*, *ANAC002* and *ATAF2*, respectively (Fig 8e). This suggests the presence of a crosstalk pathway that mediates stress tolerance between different stresses. Although these genes are common between stresses, they have different expression patterns in response to different stresses and perhaps they are involved in different gene regulatory networks. For instance, *AUR62029344*, a *ANAC072*/*RD26* orthologue, was induced in the shoots and roots of quinoa plants after salt, phosphate and drought stresses, and only downregulated in response to heat stress in both shoots and roots.

The consistent transcriptional responses between *CqNACs* (in response to abiotic stress) and those that have been reported previously for Arabidopsis or other plant species, indicate a conserved function of NACs across different species and the possible sharing of a common stress tolerance mechanism. However, we have also identified some *CqNAC*s that are stress responsive, transcriptionally, but their function have not been identified previously. These NACs may participate in a novel stress tolerance in quinoa plants.

#### 2.4.3 qRT-PCR analysis of stress responsive *CqNAC*s

To verify the response of *CqNACs* to different abiotic stresses, we conducted an additional salinity, drought and heat experiments and investigated the expression of some selected *CqNAC* by qRT-PCR. We identified the selected *CqNACs* to be DE in response to stress according to our RNAseq data and/or are orthologue to NACs functionally annotated as a stress responsive gene in other plant species. We analysed the expression of six salt stress responsive *CqNAC* genes in response to salt stress (300 mM NaCl). These genes are orthologues of Arabidopsis *ATAF1*, *ATAF2*, *ANAC072*, *ANAC078* and *ANAC083* (Fig 9a). The same trend for changes in expression were observed for all of the tested genes when comparing qRT-PCR and RNAseq (S3a Fig), which confirms their response to salt stress. Among these genes, we tested the expression of the two copies of *ANAC083* orthologues (*AUR62037478*, *AUR62043497*) and the results showed a stronger response in one of the copies (*AUR62043497*) compared to the other copy (*AUR62037478*).

**Fig 9.**
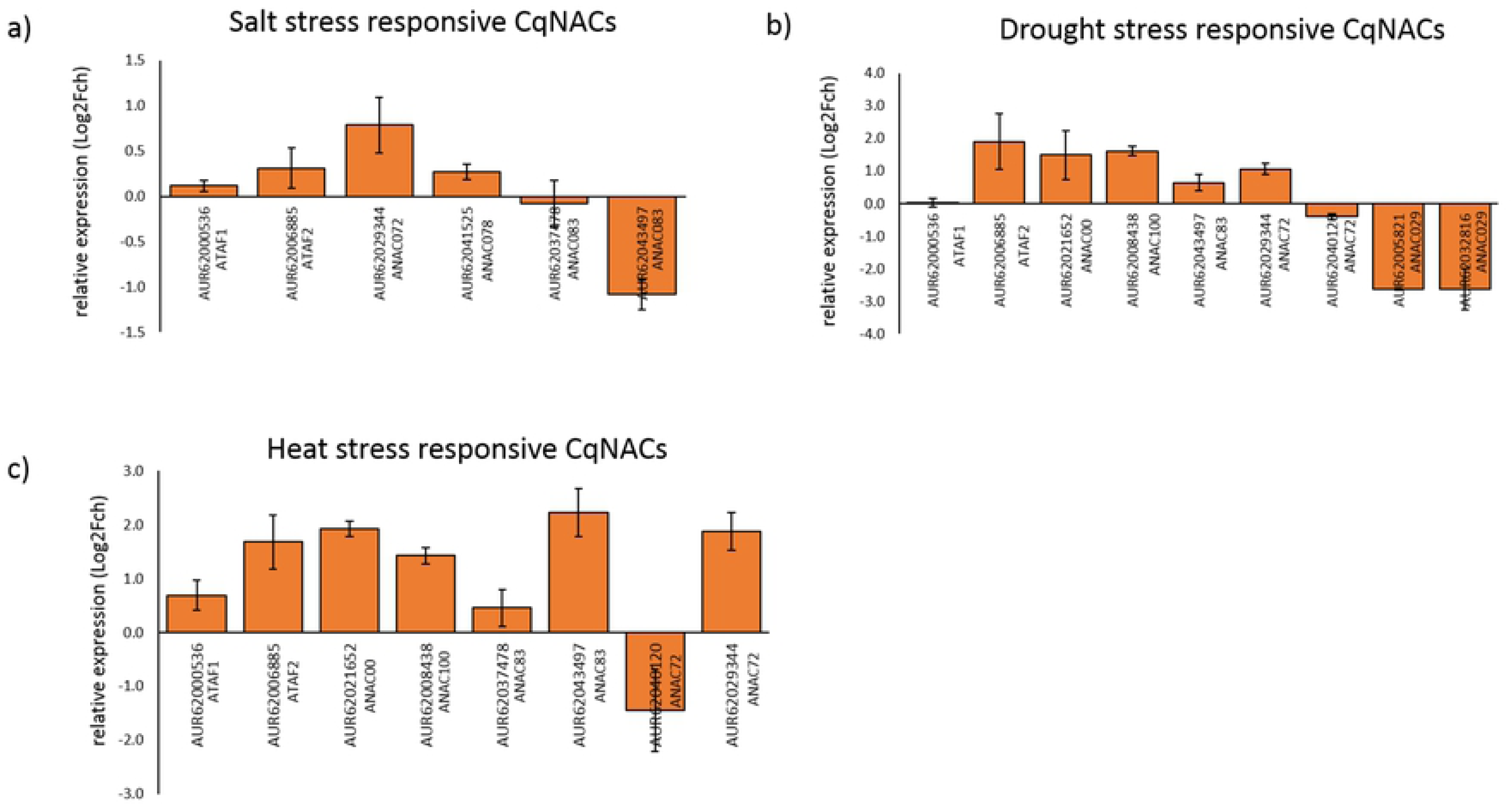
qRT-PCR analysis of some selected *CqNACs* in response to different abiotic stresses. a) Relative expression of *CqNAC*s after salt treatment relative to control. b) Relative expression of *CqNAC*s after drought treatment relative to control. a) Relative expression of *CqNAC*s after heat treatment relative to control. The expression level represents the mean of three independent biological replicates. The error bar is standard error of the mean.

In response to drought stress, we analysed the expression of nine genes (Fig 9b). The results showed that the expression pattern of some of the *CqNAC* genes in response to drought stress is consistent between RNAseq and qRT-PCR, except for four genes (*ATAF1*, *ANAC083* and the two copies of *ANAC029*), which showed a different expression pattern between RNAseq and qRT-PCR (S3b Fig). We also analysed the expression pattern between the copies of the duplicated genes, i.e., the orthologues of Arabidopsis *ANAC100*, *ANAC072* and *ANAC029*. The orthologues of Arabidopsis *ANAC100* and *ANAC072* showed a similar expression pattern between the duplicated copies, which could suggest that both of the copies have a similar function. The two copies of *ANAC072* showed an opposite expression pattern in response to drought between the two copies (RNAseq and qRT-PCR) (Fig 9b) raising the possibility that these two genes have a different function in response to drought stress in quinoa.

In response to heat stress, we analysed the expression of eight genes. The expression of all of the *CqNAC* genes in response to heat stress was consistent between the RNAseq and qRT-PCR for six of the genes. Two genes appeared to have opposite expression patterns between the RNAseq and qRT-PCR, the orthologues of Arabidopsis *ANAC083* and *ANAC072* (S3c Fig). The experiments for heat stress for RNAseq and qRT-PCR studies were performed under different conditions. The heat stress for RNAseq samples was for much shorter duration, which may explain the different expression pattern of some genes.

Among the analysed genes, there are some duplicated genes, namely the orthologues of Arabidopsis *ANAC100*, *ANAC083* and *ANAC072*. In the two duplicated genes (*ANAC100* and *ANAC083*), both copies of each gene showed a similar expression pattern in response to heat stress. This is similar to what we have seen in response to drought stress for these two genes. However, the two copies of *ANAC072* showed a different expression pattern in response to heat stress, suggesting that these two copies of *ANAC072* gene may have a different function (Fig 9c).

To conclude, the high consistency of *CqNAC* expression between the RNAseq and qRT-PCR experiments confirmed their response to abiotic stress. Moreover, by analysing the expression among the duplicated genes in response to stress, our results indicate that there is a similarity between the expression pattern and some of the duplicated genes, which indicates similar or redundant functions of these duplicated copies. However, in some duplicated genes, the expression of one copy is weaker or different (opposite) to the expression of the other copy, which may suggest different functions of the duplicated genes in response to stress. We propose that these genes should be further investigated for their role in stress responses and potentially stress tolerance.

## 3. Conclusions

In conclusion, this study provides a comprehensive identification and characterization on the transcriptional level of the NAC TFs family in quinoa. We identified 107 NAC TFs in the genome of quinoa plants. We phylogenetically and functionally classified these NACs into different phylogenetic subgroups, in alignment with previously identified NACs in Arabidopsis. This functional classification was also supported by the global expression analysis that we carried out in two tissues (shoots and roots) in response to abiotic stresses, as several NACs involved in root development were significantly differentially expressed in the root of quinoa plants. Similarly, we found several NACs responded transcriptionally to different abiotic stresses in a similar manner to previously characterized NACs in Arabidopsis and other plants species. This indicates that some of the quinoa NAC TFs have a similar function to previously identified NACs. Our results provide valuable information for further functional research of NAC TF in quinoa and their role in its adaptation to different stresses.

## 4. Materials and Methods

### 4.1 Sequence database search

We employed three approaches to identify putative NAC family genes in quinoa. In the first approach, we searched for peptides that have a protein family (Pfam) (PF02365) in the genome of quinoa in the phytozome database version12 (phytozome, http://www.phytozome.net, version 12). In the second approach, we downloaded all protein sequences of the quinoa (44,776 peptide sequences) from phytozome database (phytozome, http://www.phytozome.net, version 12), and we used these sequences as a query in the transcription factors prediction server from PTFDB. In the third approach, we used a Basic Local Alignment search tool (BLASTp); the protein sequences of the 110 NAC domain of *Arabidopsis thaliana* (Arabidopsis) were used to BLASTp against the quinoa QQ74 peptides. We selected the protein sequence with E-value of < 1e^-10^ as a candidate NAC gene, keeping only alignments that cover at least 60% of the query length (Arabidopsis sequences). We downloaded sequences of Arabidopsis and rice NAC proteins from Phytozome database version12. We considered peptide sequences that had less than 100 residues to be truncated proteins and thus they were removed from the analysis. We confirmed all of the putative NAC proteins by verifying the presence of the NAC domain with the InterProScan program (http://www.ebi.ac.uk/Tools/InterProScan/), hmmscan function of the HMMER web server (https://www.ebi.ac.uk/Tools/hmmer/search/hmmscan, HmmerWeb version 2.28.0) of NAM domain (PF02365) (33) and a conserved domain search (CDS) of the NCBI database.

We performed the inference of homologous copies of genes using a software that searches for multiple collinearity regions (MCScanX) with default parameters. Briefly, we performed between all quinoa QQ74 peptide sequences against themselves with an E-value of < 1e^-10^ and we reported a maximum of 5 best hits. Subsequently, we inferred pairwise collinear genes from collinear blocks defined by a minimum of 5 homologous sequences in conserved order. We inferred homologous genes from collinear genes between A and B sub-genomes.

The prediction of membrane-bound NACs proteins was performed using the software TMHMM (http://www.cbs.dtu.dk/services/TMHMM/, server v.2.0) (74).

### 4.2 Phylogenetic tree analysis

Multiple sequence alignment of the full length protein sequences or NAC domain sequences was performed using the tool MUSCLE (75) built in the software MEGA 7.0 (76). We constructed the unrooted phylogenetic tree using the software MEGA 7.0 (76) using the Neighbor-Joining (NJ) method with a bootstrap test value of 1000.

### 4.3 Gene structure and conserved motif

We used the gene structure display server (GSDS) program (http://gsds.cbi.pku.edu.cn/, version 2.0) (77) to display the organization of exons and introns of NAC genes by comparing cDNA with the DNA sequences of individual gene.

### 4.4 Plant material, growth conditions and stress treatment and transcriptomic analysis

We used the gene expression (RNAseq) data previously generated (27). Plant growth conditions, samples collections and stress application have been previously described (27). Briefly, RNA was isolated from roots and shoots of quinoa plants exposed to different stresses including drought, heat, salinity and low phosphate conditions. For drought and heat treatments, plants were grown in soil, for salinity and low phosphate treatments, plants were grown in hydroponics (along with their respective controls without treatment): Soil grown plants: Plants were grown in soil in a growth chamber under well-watered conditions at 20°C and 12 h daily light for three weeks and then either left without water for one week (as drought treatment) or transferred to another growth chamber with 37°C day and 32°C night temperatures (as heat treatment).

Hydroponically grown plants: After germination on agar for two weeks, seedlings were transferred to aerated tanks containing basal nutrient solution for another week before either being moved to tanks containing basal nutrient solution lacking KH_2_PO_4_ and supplemented with a compensatory amount of KCl (as low phosphate treatment) or to tanks containing basal nutrient solution supplemented with 150 mM NaCl and then increase to 300 mM NaCl after 24 h (as salinity treatment). Paired-end sequencing of 100-bp libraries was performed using an Illumina HiSeq2000. Three replicates for each treatment and each tissue were sequenced, one sample for salt treated roots was excluded as it did not pass quality control.

### 4.5 Expression analysis of *CqNAC* genes using qRT-PCR

Quinoa accession QQ74 was grown in 10 cm diameter pots with standard potting mix, under controlled conditions (day/night, 12/12 h, 22/18, 50% humidity, photon flux density 300-350 μmol m^-2^ s^-1^) in plant growth Biochambers until the 11^th^ leaf growth stage (∼26 days) before they were subjected to treatment.

Plants in all treatment groups, other than drought, were watered on the first day of treatment by filling their trays with water (or saline water in case of salt stress). Plants were also watered throughout the course of experiment to maintain pots at a water holding capacity of 60%, except for salt and drought treatments.

For heat stress, plants were moved to a chamber with the same settings as control, only the temperature was set to day/night 35/30°C. For drought stress, plants were watered up to 20-30% water holding capacity WHC (For drought plants). For salt treatment, pots were soaked in 300 mM NaCl solution for four hours.

For gene expression analysis, leaf 10 was sampled six days after the treatment, frozen in liquid nitrogen and then RNA was extracted using the Direct-Zol Plant RNA extraction kit according to the manufacturer’s instruction (Zymo Research, Germany). Quality of the RNA was assessed using gel electrophoresis and quantity using a NanoDrop. For cDNA synthesis, 1 μg of high-quality RNA was used to prepare cDNA using the SuperScript III kit according to the manufacturer’s instruction (Life Technologies).

We prepared the real time qRT-PCR reaction using the SYBR Green master mix (Applied Biosystems): cDNA 4 μL, SYBR Green master mix (2X) 5 μL, 1 μL of forward and reverse primer mix (final concentration of each primer is 0.5 μM). We incubated the reactions in the ABI PRISM 7900 HT sequence detection system (Applied Biosystems). We set the reaction using the following conditions: initial step of 95°C for 10 min followed by 40 cycles of these two steps (95°C for 15 sec, 60°C for 1 min). We used polyubiquitin 10 gene (*AUR62015654*) as the reference gene for data analysis. The primer sequences of all of the primers is provided in S7 Table. After 40 reaction cycles, we generated a melting curve by heating from 60°C to 95°C with a ramp speed of 1.9°C min^-1^ to verify amplification of the desired product.

We analysed data using the comparative Ct method as described in Kamranfar, Xue (78). Briefly, we calculated the delta Ct value (ΔCt) by normalizing each Ct value with the Ct value of the reference gene *UBQ10*. Then, the level of gene expression was expressed as the difference between an arbitrary value of 40 and the ΔCt value (a high 40-ΔCt value indicates a high gene expression level). We used a threshold of 1.5-fold change to select differentially expressed genes.

## Acknowledgements

We thank Noha Saber (KAUST, Saudi Arabia) for practical support.

## Supporting Information

**S1 Fig. Bar graph showing the number of genes in each transcription factor family in Arabidopsis and quinoa.**

Data generated using the plant transcription factor prediction tool in the plant transcription factor database (PTFDB, http://planttfdb.cbi.pku.edu.cn/, version 4.0) (32).

**S2 Fig. Distribution of NAC genes along the genome.**

a) CqNAC distribution across sub-genome A and sub-genome B. b) Number of CqNAC genes per chromosome.

**S2 Fig. Comparison of RNAseq with qRT-PCR results.**

Relative expression of *CqNACs* after treatment relative to control. a) Expression after salt treatment. b) Expression after drought treatment. c) Expression after heat treatment.

**S1 Table. TFs in quinoa predicted by PTFDB.**

List of all predicted transcription factors belonging to different families in the genome of quinoa as predicted by the plant transcription factor prediction tool in the plant transcription factor database (PTFDB) (32).

**S2 Table. Predicted NACs in quinoa.**

List of predicted NACs obtained from the three search tools including BLASTp, Pfam and PTFDB, and list of CqNACs validated by HHmer and prosite scanning tool.

**S3 Table: List of CqNACs and their homoeologes according to BLASTp and synteny analysis.**

a) List of CqNACs that has a single homoeolog in same cluster, 52 genes “26 pairs”. b) List of CqNACs that has single homoeolog in different clusters (7 pairs “14genes”). c) List of CqNACs that has more than one homolog in different cluster, (16 genes). d) List of CqNACs that has no homolog according our parameters “25 genes”.

**S4 Table: Comparison between phylogenetic subgroups of CqNACs and AtNACs.**

a) Number of Arabidopsis and quinoa NAC members in each phylogenetic subgroups. b) Function of genes in the phylogenetic subgroups 8, 16 and 18 that are absent in quinoa.

**S5 Table: Cumulative expression from RNAseq data of all of the Quinoa CqNACs in all of the treatments.**

**S6 Table: Expression values of duplicated CqNACs in control and stress conditions (salt, drought, heat and phosphate starvation).**

Expression values are based on RNAseq of three biological replicates. The significance of gene expression differences was calculated at FDR=0.05.

**S7 Table: List of primer sequences used in qRT-PCR reaction.**

## References

1. Aida M, Ishida T, Fukaki H, Fujisawa H, Tasaka M. Genes involved in organ separation in Arabidopsis: an analysis of the cup-shaped cotyledon mutant. The Plant Cell Online. 1997;9(6):841–57.

2. Souer E, van Houwelingen A, Kloos D, Mol J, Koes R. The no apical meristem gene of Petunia is required for pattern formation in embryos and flowers and is expressed at meristem and primordia boundaries. Cell. 1996;85(2):159–70.

3. Nuruzzaman M, Manimekalai R, Sharoni AM, Satoh K, Kondoh H, Ooka H, et al. Genome-wide analysis of NAC transcription factor family in rice. Gene. 2010;465(1-2):30–44.

4. Le DT, Nishiyama R, Watanabe Y, Mochida K, Yamaguchi-Shinozaki K, Shinozaki K, et al. Genome-wide survey and expression analysis of the plant-specific NAC transcription factor family in soybean during development and dehydration stress. DNA research. 2011;18(4):263–76.

5. Sablowski RW, Meyerowitz EM. A homolog of NO APICAL MERISTEM is an immediate target of the floral homeotic genes APETALA3/PISTILLATA. Cell. 1998;92(1):93–103.

6. Uauy C, Distelfeld A, Fahima T, Blechl A, Dubcovsky J. A NAC gene regulating senescence improves grain protein, zinc, and iron content in wheat. Science. 2006;314(5803):1298–301.

7. Balazadeh S, Riaño-Pachón D, Mueller-Roeber B. Transcription factors regulating leaf senescence in Arabidopsis thaliana. Plant Biology. 2008;10(s1):63–75.

8. Kim YS, Kim SG, Park JE, Park HY, Lim MH, Chua NH, et al. A membrane-bound NAC transcription factor regulates cell division in Arabidopsis. Plant Cell. 2006;18(11):3132–44.

9. Zhong R, Richardson EA, Ye Z-H. Two NAC domain transcription factors, SND1 and NST1, function redundantly in regulation of secondary wall synthesis in fibers of Arabidopsis. Planta. 2007;225(6):1603–11.

10. He XJ, Mu RL, Cao WH, Zhang ZG, Zhang JS, Chen SY. AtNAC2, a transcription factor downstream of ethylene and auxin signaling pathways, is involved in salt stress response and lateral root development. The Plant Journal. 2005;44(6):903–16.

11. Ernst HA, Nina Olsen A, Skriver K, Larsen S, Lo Leggio L. Structure of the conserved domain of ANAC, a member of the NAC family of transcription factors. EMBO reports. 2004;5(3):297–303.

12. Jensen MK, Kjaersgaard T, Nielsen MM, Galberg P, Petersen K, O’Shea C, et al. The Arabidopsis thaliana NAC transcription factor family: structure-function relationships and determinants of ANAC019 stress signalling. The Biochemical journal. 2010;426(2):183–96.

13. Olsen AN, Ernst HA, Leggio LL, Skriver K. NAC transcription factors: structurally distinct, functionally diverse. Trends in plant science. 2005;10(2):79–87.

14. Olsen AN, Ernst HA, Leggio LL, Skriver K. DNA-binding specificity and molecular functions of NAC transcription factors. Plant Science. 2005;169(4):785–97.

15. Kim SY, Kim SG, Kim YS, Seo PJ, Bae M, Yoon HK, et al. Exploring membrane-associated NAC transcription factors in Arabidopsis: implications for membrane biology in genome regulation. Nucleic Acids Research. 2007;35(1):203–13.

16. Shen H, Yin Y, Chen F, Xu Y, Dixon RA. A Bioinformatic Analysis of NAC Genes for Plant Cell Wall Development in Relation to Lignocellulosic Bioenergy Production. BioEnergy Research. 2009;2(4):217–32.

17. Christiansen MW, Holm PB, Gregersen PL. Characterization of barley (*Hordeum vulgare L.*) NAC transcription factors suggests conserved functions compared to both monocots and dicots. BMC Res Notes. 2011;4:302.

18. Yoshiyama K, Conklin PA, Huefner ND, Britt AB. Suppressor of gamma response 1 (SOG1) encodes a putative transcription factor governing multiple responses to DNA damage. Proceedings of the National Academy of Sciences. 2009;106(31):12843–8.

19. Mitsuda N, Hisabori T, Takeyasu K, Sato MH. VOZ; isolation and characterization of novel vascular plant transcription factors with a one-zinc finger from Arabidopsis thaliana. Plant and cell physiology. 2004;45(7):845–54.

20. Kolano B, McCann J, Orzechowska M, Siwinska D, Temsch E, Weiss-Schneeweiss H. Molecular and cytogenetic evidence for an allotetraploid origin of *Chenopodium quinoa* and *C. berlandieri* (Amaranthaceae). Molecular phylogenetics and evolution. 2016;100:109–23.

21. Repo-Carrasco-Valencia R, Espinoza C, Jacobsen S-E. Nutritional Value and Use of the Andean Crops Quinoa (Chenopodium quinoa) and Kañiwa (Chenopodium pallidicaule)2003. 179–89 p.

22. Isabelle Adolf V, Jacobsen S-E, Shabala S. Salt tolerance mechanisms in quinoa (Chenopodium quinoa Willd.)2013. 43–54 p.

23. Jacobsen SE, Mujica A, Jensen CR. The Resistance of Quinoa (*Chenopodium quinoaWilld.*) to Adverse Abiotic Factors. Food Reviews International. 2003;19(1-2):99–109.

24. Hariadi Y, Marandon K, Tian Y, Jacobsen SE, Shabala S. Ionic and osmotic relations in quinoa (*Chenopodium quinoa Willd.*) plants grown at various salinity levels. Journal of experimental botany. 2011;62(1):185–93.

25. Yue H, Chang X, Zhi Y, Wang L, Xing G, Song W, et al. Evolution and Identification of the WRKY Gene Family in Quinoa (*Chenopodium quinoa*). Genes. 2019;10(2).

26. Liu J, Wang R, Liu W, Zhang H, Guo Y, Wen R. Genome-Wide Characterization of Heat-Shock Protein 70s from *Chenopodium quinoa* and Expression Analyses of Cqhsp70s in Response to Drought Stress. Genes. 2018;9(2).

27. Jarvis DE, Ho YS, Lightfoot DJ, Schmöckel SM, Li B, Borm TJA, et al. The genome of *Chenopodium quinoa*. Nature. 2017;542:307.

28. Yan H, Zhang A, Ye Y, Xu B, Chen J, He X, et al. Genome-wide survey of switchgrass NACs family provides new insights into motif and structure arrangements and reveals stress-related and tissue-specific NACs. Scientific reports. 2017;7(1):3056.

29. Hu R, Qi G, Kong Y, Kong D, Gao Q, Zhou G. Comprehensive Analysis of NAC Domain Transcription Factor Gene Family in Populus trichocarpa. BMC Plant Biology. 2010;10(1):145.

30. Shang H, Li W, Zou C, Yuan Y. Analyses of the NAC Transcription Factor Gene Family in *Gossypium raimondii Ulbr*.: Chromosomal Location, Structure, Phylogeny, and Expression Patterns. Journal of Integrative Plant Biology. 2013;55(7):663–76.

31. Sun H, Hu M, Li J, Chen L, Li M, Zhang S, et al. Comprehensive analysis of NAC transcription factors uncovers their roles during fiber development and stress response in cotton. BMC Plant Biology. 2018;18(1):150.

32. Jin J, Tian F, Yang D-C, Meng Y-Q, Kong L, Luo J, et al. PlantTFDB 4.0: toward a central hub for transcription factors and regulatory interactions in plants. Nucleic Acids Research. 2017;45(D1):D1040–D5.

33. Potter SC, Luciani A, Eddy SR, Park Y, Lopez R, Finn RD. HMMER web server: 2018 update. Nucleic Acids Research. 2018;46(W1):W200–W4.

34. Ooka H, Satoh K, Doi K, Nagata T, Otomo Y, Murakami K, et al. Comprehensive analysis of NAC family genes in *Oryza sativa* and *Arabidopsis thaliana*. DNA Research 2003;10:239–47.

35. Zhang Y, Wang L. The WRKY transcription factor superfamily: its origin in eukaryotes and expansion in plants. BMC evolutionary biology. 2005;5:1.

36. Wilkins O, Nahal H, Foong J, Provart NJ, Campbell MM. Expansion and diversification of the Populus R2R3-MYB family of transcription factors. Plant physiology. 2009;149(2):981–93.

37. Li X, Duan X, Jiang H, Sun Y, Tang Y, Yuan Z, et al. Genome-wide analysis of basic/helix-loop-helix transcription factor family in rice and Arabidopsis. Plant physiology. 2006;141(4):1167–84.

38. Lijavetzky D, Carbonero P, Vicente-Carbajosa J. Genome-wide comparative phylogenetic analysis of the rice and Arabidopsis Dof gene families. BMC evolutionary biology. 2003;3:17.

39. Wang D, Guo Y, Wu C, Yang G, Li Y, Zheng C. Genome-wide analysis of CCCH zinc finger family in Arabidopsis and rice. BMC Genomics. 2008;9(1):44.

40. Hibara K, Karim MR, Takada S, Taoka K, Furutani M, Aida M, et al. Arabidopsis CUP-SHAPED COTYLEDON3 regulates postembryonic shoot meristem and organ boundary formation. Plant Cell. 2006;18(11):2946–57.

41. Vroemen CW, Mordhorst AP, Albrecht C, Kwaaitaal MACJ, de Vries SC. The CUP-SHAPED COTYLEDON3 Gene Is Required for Boundary and Shoot Meristem Formation in Arabidopsis. The Plant Cell. 2003;15(7):1563–77.

42. Zhou J, Zhong R, Ye ZH. Arabidopsis NAC domain proteins, VND1 to VND5, are transcriptional regulators of secondary wall biosynthesis in vessels. PLoS One. 2014;9(8):e105726.

43. Willemsen V, Bauch M, Bennett T, Campilho A, Wolkenfelt H, Xu J, et al. The NAC domain transcription factors FEZ and SOMBRERO control the orientation of cell division plane in Arabidopsis root stem cells. Developmental cell. 2008;15(6):913–22.

44. Tran L-SP, Nakashima K, Sakuma Y, Simpson SD, Fujita Y, Maruyama K, et al. Isolation and Functional Analysis of Arabidopsis Stress-Inducible NAC Transcription Factors That Bind to a Drought-Responsive cis-Element in the early responsive to dehydration stress 1 Promoter. The Plant Cell. 2004;16(9):2481–98.

45. Zhao J, Liu JS, Meng FN, Zhang ZZ, Long H, Lin WH, et al. ANAC005 is a membrane-associated transcription factor and regulates vascular development in Arabidopsis. Journal of integrative plant biology. 2016;58(5):442–51.

46. Kim YS, Park CM. Membrane regulation of cytokinin-mediated cell division in Arabidopsis. Plant signaling & behavior. 2007;2(1):15–6.

47. Rahman H, Ramanathan V, Nallathambi J, Duraialagaraja S, Muthurajan R. Over-expression of a NAC 67 transcription factor from finger millet (*Eleusine coracana L.*) confers tolerance against salinity and drought stress in rice. BMC biotechnology. 2016;16 Suppl 1:35.

48. Jung JH, Park CM. Auxin modulation of salt stress signaling in Arabidopsis seed germination. Plant signaling & behavior. 2011;6(8):1198–200.

49. Prince VE, Pickett FB. Splitting pairs: the diverging fates of duplicated genes. Nature reviews Genetics. 2002;3(11):827–37.

50. Liu M, Ma Z, Sun W, Huang L, Wu Q, Tang Z, et al. Genome-wide analysis of the NAC transcription factor family in Tartary buckwheat (*Fagopyrum tataricum*). BMC Genomics. 2019;20(1):113.

51. Shahnejat-Bushehri S, Tarkowska D, Sakuraba Y, Balazadeh S. Arabidopsis NAC transcription factor JUB1 regulates GA/BR metabolism and signalling. Nature plants. 2016;2.

52. Thirumalaikumar VP, Devkar V, Mehterov N, Ali S, Ozgur R, Turkan I, et al. NAC transcription factor JUNGBRUNNEN1 enhances drought tolerance in tomato. Plant Biotechnology Journal. 2018;16(2):354–66.

53. Lee S, Lee HJ, Huh SU, Paek KH, Ha JH, Park CM. The Arabidopsis NAC transcription factor NTL4 participates in a positive feedback loop that induces programmed cell death under heat stress conditions. Plant science : an international journal of experimental plant biology. 2014;227:76–83.

54. He XJ, Mu RL, Cao WH, Zhang ZG, Zhang JS, Chen SY. AtNAC2, a transcription factor downstream of ethylene and auxin signaling pathways, is involved in salt stress response and lateral root development. The Plant journal : for cell and molecular biology. 2005;44(6):903–16.

55. De Zelicourt A, Diet A, Marion J, Laffont C, Ariel F, Moison M, et al. Dual involvement of a Medicago truncatula NAC transcription factor in root abiotic stress response and symbiotic nodule senescence. The Plant journal : for cell and molecular biology. 2012;70(2):220–30.

56. Zhong R, Ye Z-H. The Arabidopsis NAC transcription factor NST2 functions together with SND1 and NST1 to regulate secondary wall biosynthesis in fibers of inflorescence stems. Plant signaling & behavior. 2015;10(2):e989746-e.

57. Nuruzzaman M, Sharoni AM, Kikuchi S. Roles of NAC transcription factors in the regulation of biotic and abiotic stress responses in plants. Frontiers in Microbiology. 2013;4:248.

58. Golldack D, Lüking I, Yang O. Plant tolerance to drought and salinity: stress regulating transcription factors and their functional significance in the cellular transcriptional network. Plant cell reports. 2011;30(8):1383–91.

59. Chen X, Wang Y, Lv B, Li J, Luo L, Lu S, et al. The NAC family transcription factor OsNAP confers abiotic stress response through the ABA pathway. Plant & cell physiology. 2014;55(3):604–19.

60. Huang Q, Wang Y, Li B, Chang J, Chen M, Li K, et al. TaNAC29, a NAC transcription factor from wheat, enhances salt and drought tolerance in transgenic Arabidopsis. BMC Plant Biol. 2015;15:268.

61. Yang SD, Seo PJ, Yoon HK, Park CM. The Arabidopsis NAC transcription factor VNI2 integrates abscisic acid signals into leaf senescence via the COR/RD genes. The Plant cell. 2011;23(6):2155–68.

62. Seok HY, Woo DH, Nguyen LV, Tran HT, Tarte VN, Mehdi SM, et al. Arabidopsis AtNAP functions as a negative regulator via repression of AREB1 in salt stress response. Planta. 2017;245(2):329–41.

63. Tran LS, Nakashima K, Sakuma Y, Simpson SD, Fujita Y, Maruyama K, et al. Isolation and functional analysis of Arabidopsis stress-inducible NAC transcription factors that bind to a drought-responsive cis-element in the early responsive to dehydration stress 1 promoter. Plant Cell. 2004;16(9):2481–98.

64. Delessert C, Kazan K, Wilson IW, Van Der Straeten D, Manners J, Dennis ES, et al. The transcription factor ATAF2 represses the expression of pathogenesis-related genes in Arabidopsis. The Plant journal : for cell and molecular biology. 2005;43(5):745–57.

65. Seki M, Narusaka M, Ishida J, Nanjo T, Fujita M, Oono Y, et al. Monitoring the expression profiles of 7000 Arabidopsis genes under drought, cold and high-salinity stresses using a full-length cDNA microarray. The Plant journal : for cell and molecular biology. 2002;31(3):279–92.

66. Wu Y, Deng Z, Lai J, Zhang Y, Yang C, Yin B, et al. Dual function of Arabidopsis ATAF1 in abiotic and biotic stress responses. Cell research. 2009;19(11):1279–90.

67. Shahnejat-Bushehri S, Mueller-Roeber B, Balazadeh S. Arabidopsis NAC transcription factor JUNGBRUNNEN1 affects thermomemory-associated genes and enhances heat stress tolerance in primed and unprimed conditions. Plant Signaling & Behavior. 2012;7(12):1518–21.

68. Guan Q, Yue X, Zeng H, Zhu J. The protein phosphatase RCF2 and its interacting partner NAC019 are critical for heat stress-responsive gene regulation and thermotolerance in Arabidopsis. Plant Cell. 2014;26(1):438–53.

69. Fang Y, Liao K, Du H, Xu Y, Song H, Li X, et al. A stress-responsive NAC transcription factor SNAC3 confers heat and drought tolerance through modulation of reactive oxygen species in rice. Journal of experimental botany. 2015;66(21):6803–17.

70. Guo W, Zhang J, Zhang N, Xin M, Peng H, Hu Z, et al. The Wheat NAC Transcription Factor TaNAC2L Is Regulated at the Transcriptional and Post-Translational Levels and Promotes Heat Stress Tolerance in Transgenic Arabidopsis. PLoS One. 2015;10(8):e0135667.

71. Morcuende R, Bari R, Gibon Y, Zheng W, Pant BD, Blasing O, et al. Genome-wide reprogramming of metabolism and regulatory networks of Arabidopsis in response to phosphorus. Plant, cell & environment. 2007;30(1):85–112.

72. Misson J, Raghothama KG, Jain A, Jouhet J, Block MA, Bligny R, et al. A genome-wide transcriptional analysis using *Arabidopsis thaliana* Affymetrix gene chips determined plant responses to phosphate deprivation. Proceedings of the National Academy of Sciences of the United States of America. 2005;102(33):11934–9.

73. Nilsson L, Müller R, Nielsen TH. Dissecting the plant transcriptome and the regulatory responses to phosphate deprivation. Physiologia Plantarum. 2010;139(2):129–43.

74. Krogh A, Larsson B, von Heijne G, Sonnhammer EL. Predicting transmembrane protein topology with a hidden Markov model: application to complete genomes. Journal of molecular biology. 2001;305(3):567–80.

75. Edgar RC. MUSCLE: multiple sequence alignment with high accuracy and high throughput. Nucleic Acids Research. 2004;32(5):1792–7.

76. Kumar S, Stecher G, Tamura K. MEGA7: Molecular Evolutionary Genetics Analysis Version 7.0 for Bigger Datasets. Molecular biology and evolution. 2016;33(7):1870–4.

77. Hu B, Jin J, Guo A-Y, Zhang H, Luo J, Gao G. GSDS 2.0: an upgraded gene feature visualization server. Bioinformatics. 2015;31(8):1296–7.

78. Kamranfar I, Xue G-P, Tohge T, Sedaghatmehr M, Fernie AR, Balazadeh S, et al. Transcription factor RD26 is a key regulator of metabolic reprogramming during dark-induced senescence. New Phytologist. 2018;218(4):1543–57.

